# Self-organized patterning of cell morphology via mechanosensitive feedback

**DOI:** 10.1101/2020.04.16.044883

**Authors:** Natalie A. Dye, Marko Popovic, K. Venkatesan Iyer, Suzanne Eaton, Frank Julicher

## Abstract

Tissue organization is often characterized by specific patterns of cell morphology. How such patterns emerge in developing tissues is a fundamental open question. Here, we investigate the emergence of tissue-scale patterns of cell shape and mechanical tissue stress in the *Drosophila* wing imaginal disc during larval development. Using quantitative analysis of the cellular dynamics, we reveal a pattern of radially oriented cell rearrangements that is coupled to the buildup of tangential cell elongation. Developing a laser ablation method, we map tissue stresses and extract key parameters of tissue mechanics. We present a continuum theory showing that this pattern of cell morphology and tissue stress can arise via self-organization of a mechanical feedback that couples cell polarity to active cell rearrangements. The predictions of this model are supported by knockdown of MyoVI, a component of mechanosensitive feedback. Our work reveals a mechanism for the emergence of cellular patterns in morphogenesis.

## INTRODUCTION

During morphogenesis, tissues with complex morphologies are formed from the collective interplay of many cells. This process involves spatial patterns of signaling activity, and previous work has discovered mechanisms for generating tissue-scale patterns of activity in signaling pathways such as Hedgehog, TGF*β*, and Wnt (Green & Sharpe, 2015). In addition, patterns of cellular morphology arise during morphogenesis. Such patterns can be important for ensuring the function of the resulting tissue. For example, the compound eye of *Drosophila* consists of hundreds of ommatidia organized in a precise hexagonal array that is required to fully sample the visual field (Kumar, 2012). Patterns of cellular morphology that arise during morphogenesis can also guide the morphogenetic processes itself. For example, spatial patterns of cell morphology emerge during growth of the *Drosophila* larval imaginal discs, which are precursors of adult tissues (Aegerter-Wilmsen et al., 2010; Condic et al., 1991; LeGoff et al., 2013; Mao et al., 2013). These patterns have been proposed to be involved in the eversion process, during which these flattened epithelial sacs turn themselves inside out when the animal transitions from larva to pupa (Condic et al., 1991). While extensive work has studied the emergence of biochemical signaling patterns, how patterns of cellular morphology arise during tissue development is poorly understood.

Here, we investigate tissue-scale patterning of cell morphology in the *Drosophila* larval wing imaginal disc, which has a geometry that is ideal for studying spatial patterns of epithelial cell morphology. We focus specifically on the cell shape patterns in the central “pouch” region, which is the precursor of the adult wing blade. To a good approximation, this region is planar and ellipsoidal. Cells near the center have smaller cell areas and are more isotropic in shape, whereas cells near the periphery have larger cell areas and are elongated tangentially (Aegerter-Wilmsen et al., 2010; LeGoff et al., 2013; Mao et al., 2013). Cell shape has been correlated with mechanical stress: tangentially oriented bonds of elongated cells in the periphery are under higher tension than radially oriented bonds (LeGoff et al., 2013).

It has been previously proposed that this pattern of cell morphology in the wing pouch could stem from differential proliferation: if the center grows faster than the rest, the resulting area pressure could stretch peripheral cells tangentially (Mao et al., 2013). Indeed, there is evidence to suggest that cells divide slightly faster closer to the center during very early stages (before 80hr after egg laying, AEL). It was suggested that this early growth differential is sufficient to account for the persistence of the cell morphology pattern through the remaining ~40hrs of development. However, it has since been shown that cell rearrangements occur (Dye et al., 2017; Heller et al., 2016), which could relax stress patterns once growth has become uniform. Furthermore, stress patterns may even relax during homogeneous growth in the absence of cell rearrangements (Ranft et al., 2010). Thus, it remains unclear how cell morphology patterns generated early by differential growth could be maintained through later stages, and alternative mechanisms for the establishment of these patterns must be considered.

Here, we measure the spatial patterns of cell morphology, cell divisions, and cell rearrangements during the middle of the third larval instar (starting at 96hr AEL). We quantify the pattern of tangential cell elongation and show that it becomes stronger over time, even though growth is spatially uniform and cell rearrangements are frequent. Strikingly, this change in tangential cell elongation is coupled to a radially biased pattern of cell neighbor exchanges. Using a physical model of tissue dynamics, we show that active patterning of radial cell neighbor exchanges can account for the observed morphology patterns in the absence of differential growth. Lastly, using a combination of experiment and theory, we provide evidence that this active patterning is self-organized by mechanosensitive feedback.

## RESULTS

### Cell morphology patterns can persist and strengthen in the absence of differential growth

Cell morphology patterns in the wing disc have been previously analyzed using static images (Aegerter-Wilmsen et al., 2010; LeGoff et al., 2013; Mao et al., 2013). However, relating cell morphology patterns to patterns of growth, cell divisions and cell rearrangements requires dynamic data. We therefore performed long-term timelapse imaging of growing explanted wing discs using our previously described methods (Dye et al., 2017), starting at 96 *hr* AEL and continuing for ~13 *hr* of imaging. We used E-cadherin-GFP as an apical junction marker (Fig 1A).

**Figure 1:**
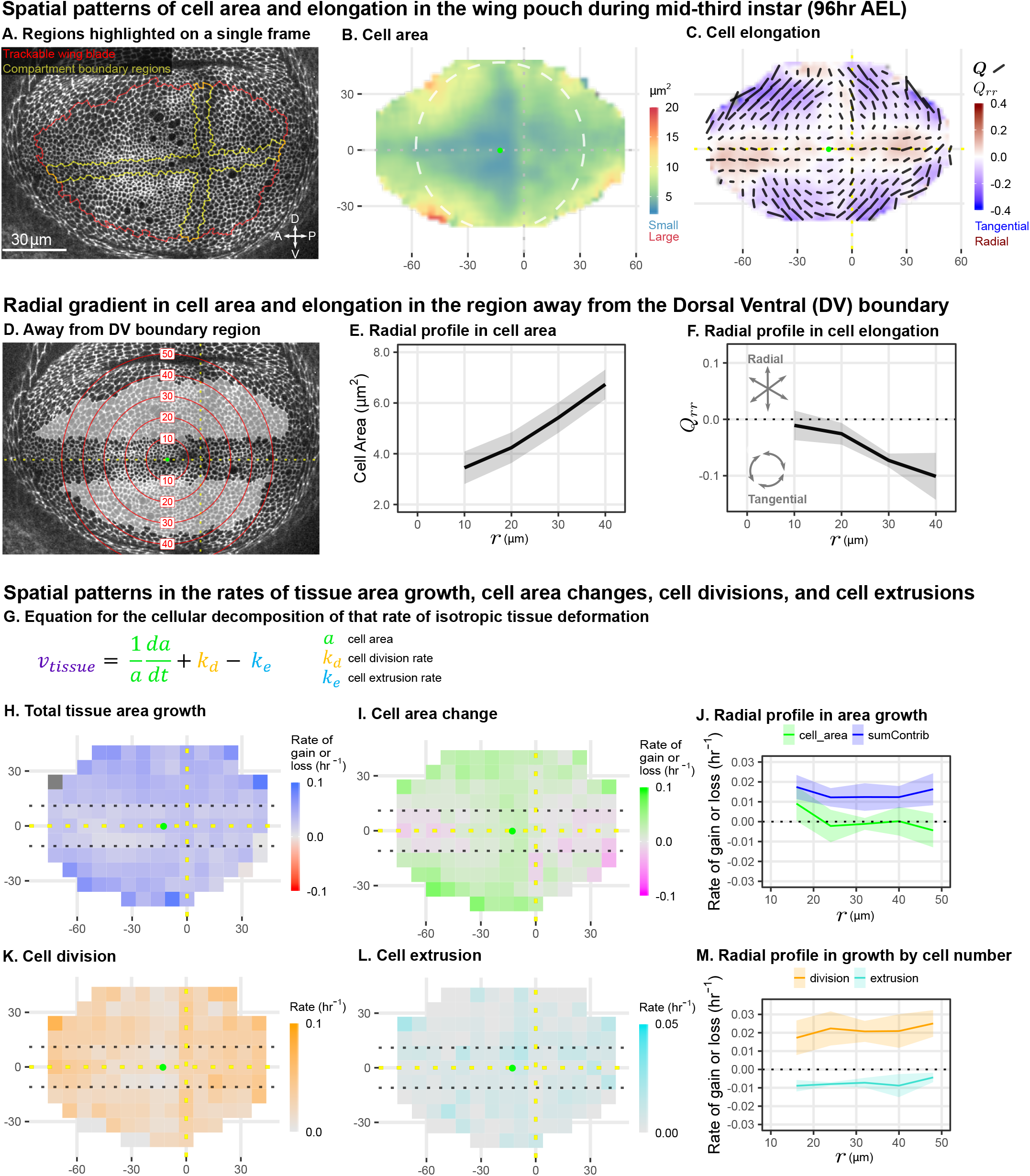
Cell morphology patterns can persist and strengthen in the absence of differential growth. (A) Ecadherin-GFP-expressing wing disc growing in *ex vivo* culture (explanted at 96 *hr* AEL). We analyze apical cell morphology in the proliferating disc proper layer in a 2D projection, after correcting for local tissue curvature (see Methods and Fig S1A). Red outline indicates the presumptive “blade” region; yellow outline indicates cells of the compartment boundaries, which are used to align different movies. Dorsal is up, and anterior is left. (B-C) Spatial maps of cell area (B) or cell elongation (C) were generated by averaging across the middle timepoints of five movies. Axis labels indicate the distance in *μm* from the AP boundary (along X) or DV boundary (along Y). Dotted lines indicate the compartment boundaries. The calculated center of symmetry is represented by a green dot. In (B), the white dashed circle is added to highlight the near-circular symmetry. In (C), the bars represent the cell elongation tensor **Q**, where the length of the bar is proportional to the magnitude and the angle indicates its orientation. In addition, the radial component of cell elongation is presented in color. More information about this cell elongation pattern and how we define the center point is included in Fig S1. (D) The blade cells that are tangentially elongated and lying well outside the DV boundary region are shaded in grey on a single image of the timelapse. Red circles indicate the radial binning used in (E-F), where the numbers indicate the radius (in *μm*). The region around the DV boundary was separately analyzed in Fig S2. For the region depicted in (D), the radial gradient in average cell area (E) and radial cell elongation *Q_rr_* (F) were calculated. Solid lines indicate the average over all five movies in the middle time window. The shaded region indicates the standard deviation. Data showing how these radial gradients change over time during imaging are shown in Fig S3. (G) Decomposition of isotropic tissue deformation into cellular contributions. (H-I, K-L) Maps of the total tissue area growth (H) and its contributions from cell area change (I), cell divisions (K), and cell extrusions (L) do not show pronounced spatial patterns. (J, M) The radial profile of tissue growth and its cellular contributions, analyzed in radial bins. sumContrib in blue is the sum of all contributions, corresponding to total tissue deformation.

To quantify morphology, we averaged apical cell area and cell elongation locally in space, using data from 5 wing discs, and in time using ~2 *hr* intervals. We present the spatial patterns calculated for the middle timepoint in Fig 1B-C. Cell area is represented as a color code. Cell elongation is characterized by a tensor, which defines an axis and a strength of elongation and is represented by bars in Fig 1C (see Methods, Fig S1B) (Etournay et al., 2015; Merkel et al., 2017). To quantify the radial symmetry of this pattern, we first determined the center of symmetry (Fig S1 and methods). We then introduce a polar coordinate system at the center with the radial coordinate *r* and present the radial projection *Q_rr_* of cell elongation tensor as a color code in Fig 1C (see also Fig S1). This figure highlights the pattern of tangential cell elongation, with cells elongating on average perpendicular to the radial axis (blue in Fig 1C). It also reveals that this pattern is interrupted around the Dorsal-Ventral (DV) boundary, where cells are elongated parallel to this boundary (red in Fig 1C). We quantify this region separately (Fig S2) and exclude it from our analysis of the circular patterns (Fig 1D-F). The Anterior-Posterior (AP) boundary also affects cell morphology (Landsberg et al., 2009), but the effect is weaker and more variable at this stage; thus, we do not quantify it separately.

The spatial maps of cell morphology reveal that both cell area and cell elongation magnitude are largest at the periphery and decrease toward the center (Fig 1B-C). We quantified this radial gradient in cell area and observe that cell area ranges from ~3 – 7*μm*^2^ when moving toward the periphery (Fig 1E). We also observe a radial gradient in cell elongation starting from *Q_rr_* ≈ 0 at *r* = 10 *μm* and extending to about *Q_rr_* ≈ −0.1 at *r* = 40 *μm* (Fig 1F). The negative value corresponds to tangential elongation (Fig 1C). When evaluated over the timelapse, we find that these radial gradients grow slightly more pronounced over time (Fig S3C-D).

As previously proposed, differential growth can generate such patterns of cell elongation (Mao et al., 2013). However, indirect metrics of tissue growth *in vivo* do not indicate that differential growth still occurs at this later stage (Mao et al., 2013). We directly measured the spatial pattern of growth during timelapse. Cell division rate has been previously used as an indicator of growth; however, tissue growth actually results from a combination of cell division, cell area changes, and cell extrusions (Fig 1G). Thus, we evaluated the spatial pattern (Fig 1H, I, K, L) and radial profiles (Fig 1J, M) of total tissue growth and its cellular contributions. These data show that tissue area growth, as well as cell division rate, are to a good approximation independent of the distance to the center.

In summary, we quantified cell morphology patterns in mid-third instar wing explants during live imaging. We confirm previous static observations of the pattern and further identify a region around the DV boundary with a morphological pattern that differs from the rest of the wing pouch. We quantify the radial gradients in cell area and cell elongation existing outside of the DV boundary region and show that they strengthen in time in the absence of differential growth, raising the question of what mechanism underlies the persistence of these cell morphology patterns during mid to late stages of wing growth.

### Radially-oriented cell rearrangements balance tangential cell elongation

To directly relate the observed cell morphology patterns with cell rearrangements, we next analyzed the spatial patterns in cellular contributions to tissue shear (Fig 2A). Tissue shear can be decomposed into contributions from cell divisions, cell elongation changes, T1 transitions, and so-called correlation effects (Etournay et al., 2015; Merkel et al., 2017). Here, correlation effects result mainly from correlated fluctuations in cell elongation and cell rotation (see Theory Supplement part III and (Merkel et al., 2017)).

**Figure 2:**
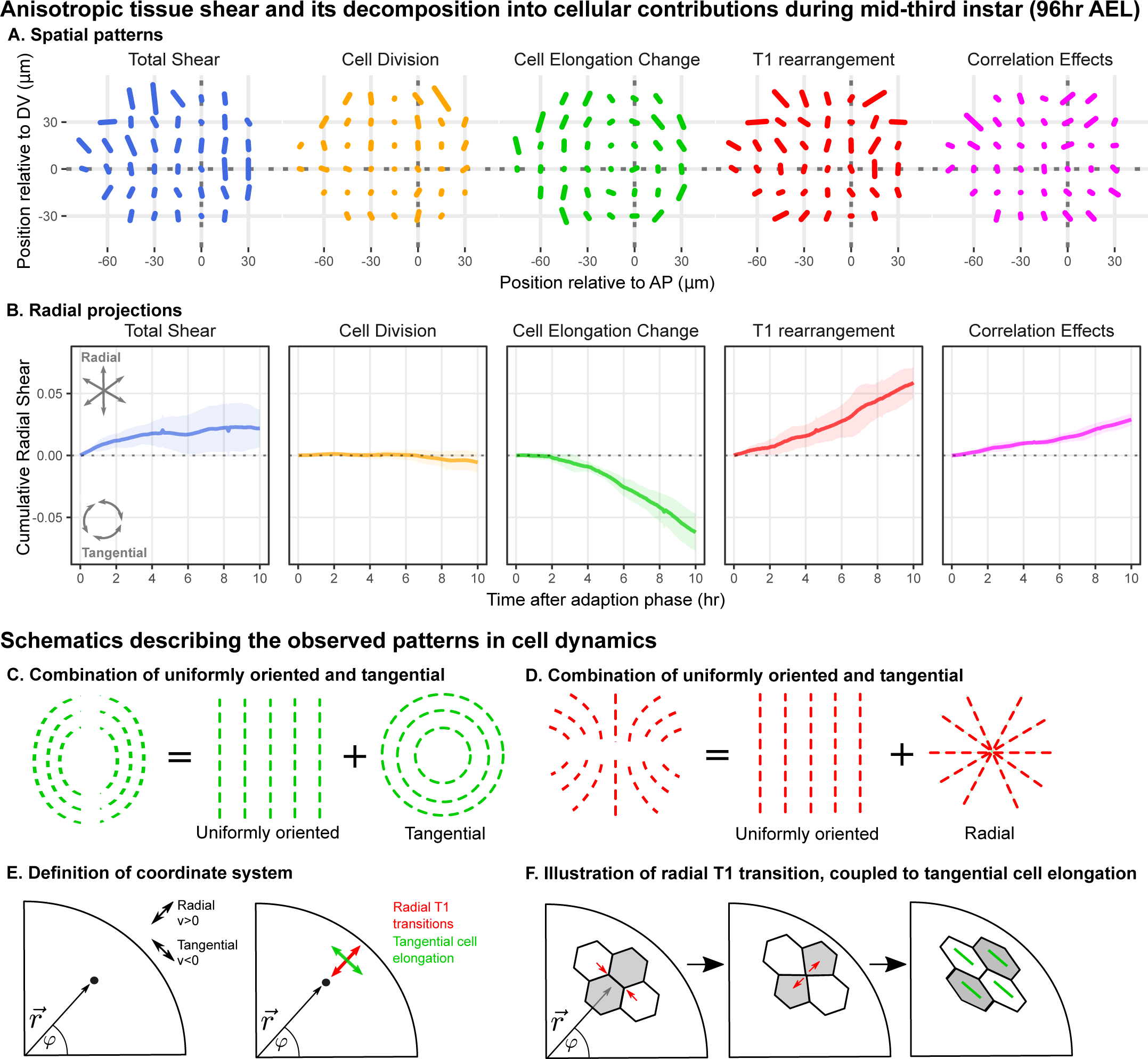
Radially-oriented cell rearrangements balance tangential cell elongation. (A) Cumulative tissue deformation and its cellular contributions, measured in a grid centered on the compartment boundaries (dotted lines) and averaged over all 5 movies. Bars represent nematic tensors, where the length is proportional to the magnitude of deformation and the angle indicates its orientation. The contribution from cell extrusion is small and thus not shown. (B) The radial projection of cumulative tissue deformation and its cellular contributions are plotted as a function of time after the first 2 *hr* of adaption to culture (Dye et al., 2017). Solid lines indicate the average over all five movies; shading indicates the standard deviation. (C) Schematics indicating how a uniformly oriented pattern would combine with a tangential pattern to produce a pattern resembling that of the cell elongation change (left) or with a radial pattern to produce a pattern resembling that of the T1 transitions (right). (E-F) Illustrations demonstrating radially-oriented T1 transitions coupled to tangential cell elongation. For simplicity, we diagram only the posterior-dorsal quadrant, but the pattern is radially symmetric. In (E), we define the radial coordinate system, where velocity in the radial direction is positive and that in the negative direction is negative. Patterns in A indicate that T1 transitions are biased to grow new bonds in the radial direction, and cell elongation is biased to increase tangentially. (F) In a radially oriented T1 transition, cells preferentially shrink tangentially oriented bonds and grow new bonds in the radial direction. When oriented in this direction, T1 transitions do not dissipate tangential elongation but increase it (green bars in each cell represent cell elongation).

We find that the spatial patterns of tissue growth and its cellular contributions exhibit overall anisotropies perpendicular to the DV boundary, as reported previously (Dye et al., 2017). In addition, the patterns of cell elongation changes and T1 transitions can be described as a superposition of a uniformly oriented pattern and an approximately radial or tangential pattern (Fig 2C-D). To determine the magnitude of the radial or tangential patterns in all quantities, we quantified their average radial projections as cumulative plots over time (Fig 2B). The radial component of tissue shear is small, and cell divisions do not contribute to radial tissue shear. In contrast, we observe a pronounced buildup of a tangential pattern of cell elongation accompanied by a radial pattern of T1 transitions and of correlation effects.

As shown above (Fig 1J), tissue area growth does not have a radial gradient and thus does not contribute to the increase in tangential cell elongation that we observe at this time (Fig 2A-B, Fig S3C-D). Furthermore, we observe numerous T1 transitions (on average 1.0 *cell*^−1^ *hr*^−1^), and their radially-biased orientation increases rather than relaxes tangential cell elongation (Fig 2E-F). Thus, we are not observing the relaxation of a pattern of cell elongation caused by early differential growth. Rather, our data support a model whereby a radially patterned morphogenetic cue actively biases the direction of T1 transitions and consequently the complementary pattern of cell shape changes.

### Polarity-driven cell rearrangements can create the observed cell morphology patterns in the wing disc

We next apply a biophysical model to determine whether radially patterned T1 transitions could account for the observed cell morphology patterns in the wing disc. We propose that the wing disc has a patterning cue that biases T1 transition orientation. This cue could be represented as a cell polarity that is nematic in nature, having a magnitude and an orientation axis, which is the same symmetry as that of an elongated cell. To capture the mechanics and dynamics of the epithelium, we adapt a physical model of the tissue material properties that takes into account the interplay of T1 transitions, cell shape changes, and tissue shear in a continuum description (Etournay et al., 2015; Popović et al., 2016). Such a model can predict the patterns of cell morphology that arise from patterned T1 transitions.

In our model, we consider the spatial patterns of tissue shear rate 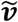, the patterns of cell shape ***Q***, and the patterns of cell rearrangements ***R***, which have nematic symmetry and are represented by traceless symmetric tensors that describe the magnitude and orientation. Tissue shear results from a combination of cell shape changes and rearrangements according to:

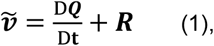

where D***Q***/D*t* is a co-rotational time derivative of ***Q***, and the shear due to rearrangements ***R*** includes contributions from T1 transitions, cell divisions and extrusions, and also correlation effects. Eq 1 follows from geometric considerations (Etournay et al., 2015; Merkel et al., 2017).

Tissue material properties are described by constitutive equations for the tissue shear stress 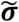 and the shear due to cell rearrangements ***R*** (Fig 3A-B):

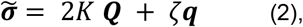

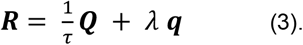

**Figure 3:**
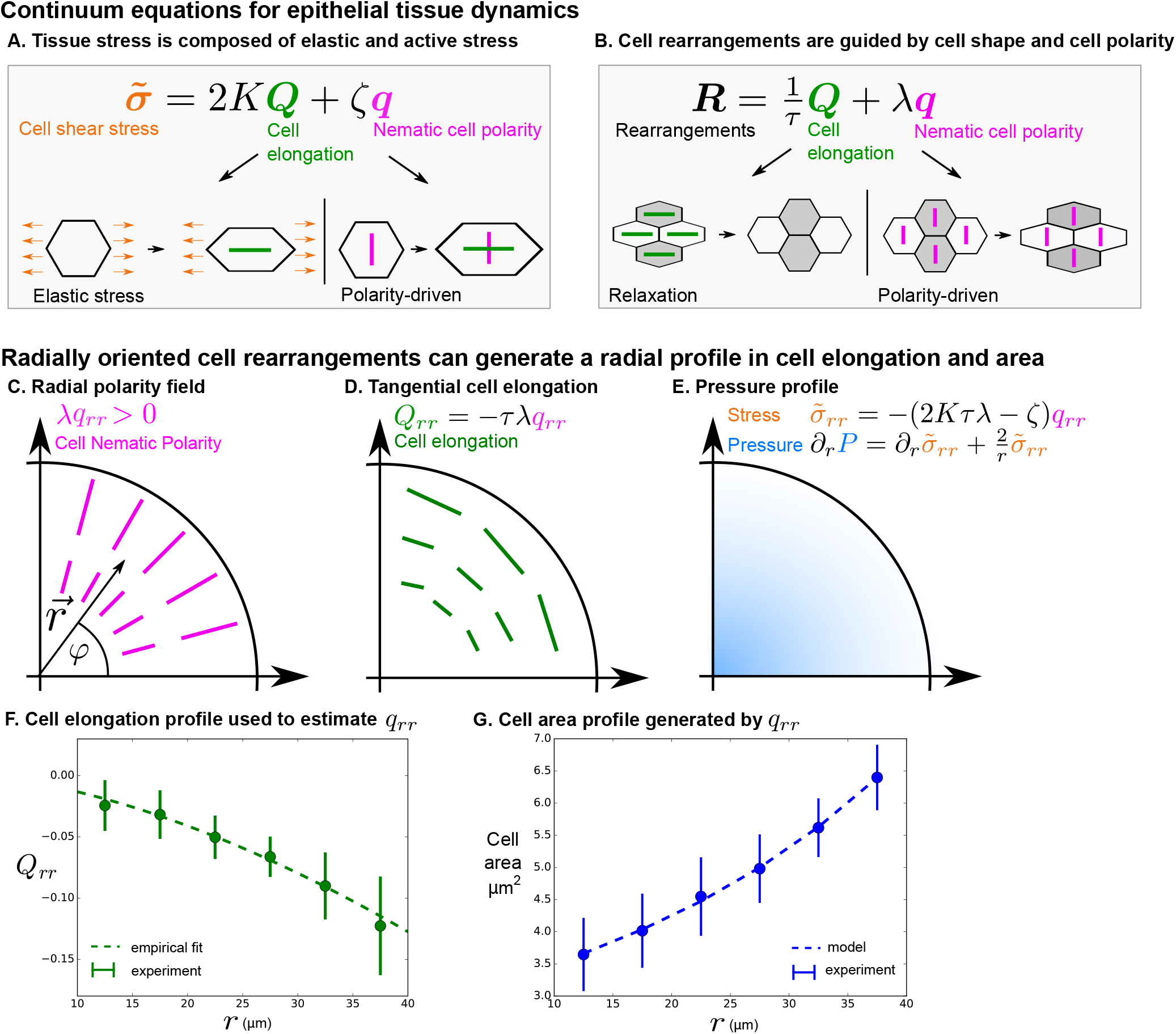
Polarity-driven cell rearrangements can create the observed cell morphology patterns. (A) Stress is a combination of elastic and polarity-driven stresses. Similarly, in (B), cell rearrangements occur to relax a stretched cell shape or to respond to an internal nematic cell polarity cue. In the cartoons in (A)-(B), we chose to depict scenarios where *ζ* and *λ* are > 0. (C-G) We apply this model to a radially symmetric tissue at steady state to approximate the wing disc. If we impose a radial polarity field (C), cell elongation is oriented in the opposite direction, according to the equation in (D) when *R_rr_* = 0 (steady state). (E) Considering force balance in the tissue, our model also predicts a pressure profile with higher pressure in the center. (F) To fit experimental data in the wing disc, we estimate the radial profile of *q_rr_* from Eq 4 by measuring cell elongation as a function of *r* in the last (~5 *hr*) of the timelapse. We solve for *q_rr_* by making an empirical fit to this cell elongation data (see Theory Supplement part IB). (G) The cell area distribution we observe in the wing disc is consistent with the pressure profile predicted by our model (E and Theory Supplement part IB).

Here, the shear stress tensor 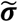 is the sum of the elastic stress associated with cell elongation ***Q*** and the active stress associated with the cell polarity cue ***q*** (Fig 3A). The elastic stress is characterized by the shear elastic modulus *K*. The cell polarity cue is provided by the nematic tensor ***q***, and the coefficient *ζ* describes the anisotropic stress associated with the polarity cue. The shear due to cell rearrangements ***R*** given by Eq 3 is driven in part by shear stress, and therefore depends on the cell elongation ***Q***, and in part by active processes that are oriented by the nematic cell polarity ***q*** (Fig 3B). *τ* is a relaxation time for cell rearrangements, and *λ* is the rate of cell rearrangements driven by polarity.

To discuss the wing disc, we consider a radially symmetric geometry and average the oriented quantities after projection onto the radial axis. Radial tissue shear is small compared to that associated with cell shape changes and T1 transitions during our observed time window (Fig 2B). We therefore consider a steady state with 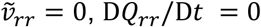, and *R_rr_* = 0. In this case, cell elongation becomes:

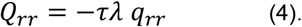

Thus, we find that the steady state cell elongation pattern is a result of cell rearrangements that are oriented by the cell polarity cue ***q*** (Fig 3C-D). Note that our data show that the wing disc is not exactly at steady state: cells slowly change their shape and rearrange radially (Fig 2A-B). However, as we show in Theory Supplement part I, Eq. 4 holds to a good approximation.

Can the radial pattern of T1 transitions defined by ***q*** also explain the observed radial profile of cell area (Fig 1B, E)? To answer this question, we then considered force balances in the tissue. We consider tissue area pressure

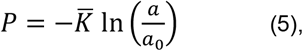

where *P* is the difference in pressure from a reference value, 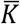 is tissue area compressibility, *a* is the average cell area, and *a*_0_ is a reference cell area. As pressure increases, cell area decreases. To calculate the cell area profile, we again approximate the wing pouch as a radially symmetric disc. In the radially symmetric geometry, force balance can be expressed as:

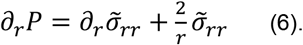

A radial profile of pressure determined from this equation implies a radial pattern of cell area via Eq 5 (see Fig 3C-E and Theory Supplement part I). To fit this equation to our data, we first estimate the profile of *q_rr_* by measuring the radial profile of cell elongation using Eq 4 (Fig 3F). Then, we solve for the cell area profile using Eqs 2, 5, and 6 (Fig 3G and Theory Supplement part I). We obtain a fit that accounts for the observed pattern of cell area.

From this analysis, we conclude that the cell morphology patterns observed in the wing disc could be generated by radially-biased cell rearrangements. Next, we test whether the stress profile predicted by the model (Eq 2) exists in the tissue, and we measure key mechanical parameters of the model. Later, we address the potential molecular origin of the cell polarity cue orienting the cell rearrangements.

### Circular laser ablation reveals patterns of tissue stress

Our model predicts a stress pattern in the wing disc that results from active processes that are radially oriented by a cell polarity cue. To compare this prediction to experiment we infer tissue stress using laser ablation. Tissue stress has been estimated previously by laser ablation techniques that are based on determining the initial retraction velocity (Bonnet et al., 2012; Etournay et al., 2015; Farhadifar et al., 2007; LeGoff et al., 2013; Mao et al., 2013; Shivakumar & Lenne, 2016). However, to compare theory and experiment, ideally one should measure quantities that are well-captured by the model. Therefore, instead of using initial retraction velocity, we perform circular cuts and analyze the final, relaxed position of the inner and outer elliptical contours of tissue formed by the cut (Fig 1A, Theory Supplement part II). From the size and the anisotropy of the cut, we can infer anisotropic and isotropic tissues stress, normalized by the respective elastic constants, as well as the ratio of elastic constants (see Theory Supplement part II). Furthermore, we can also infer the existence of polarity-driven stress. We name this method ESCA (<u>E</u>lliptical <u>S</u>hape after <u>C</u>ircular <u>A</u>blation).

We perform local measurements of tissue mechanics by cutting the tissue in the smallest possible circle that would still allow us to measure the shape of the inner piece left by the cut (radius = 7 *μm*, encircling ~ 5 – 15 cells, Fig 4A). To relate measurements of tissue stress to cell elongation, we calculate the average cell elongation in the ablated region before it is cut (Fig 4A). In Fig 4B, we present the measured cell elongation and shear stress tensors at the position of ablation. We find that local average cell elongation correlates well with the direction of shear stress. Also, we observe that the cells in the band around the DV boundary have different mechanical properties than elsewhere in the tissue. Near the DV boundary, cells elongate less than outside this region for comparable amounts of stress (Fig 4B and Fig S4B, E). The ratio of elastic constants in this region is also smaller: near the DV boundary 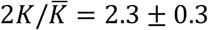, whereas outside this region, 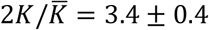 (see also Fig S4C, F). We focus hereafter on the radial patterns of elongation and stress outside of the DV boundary region.

**Figure 4:**
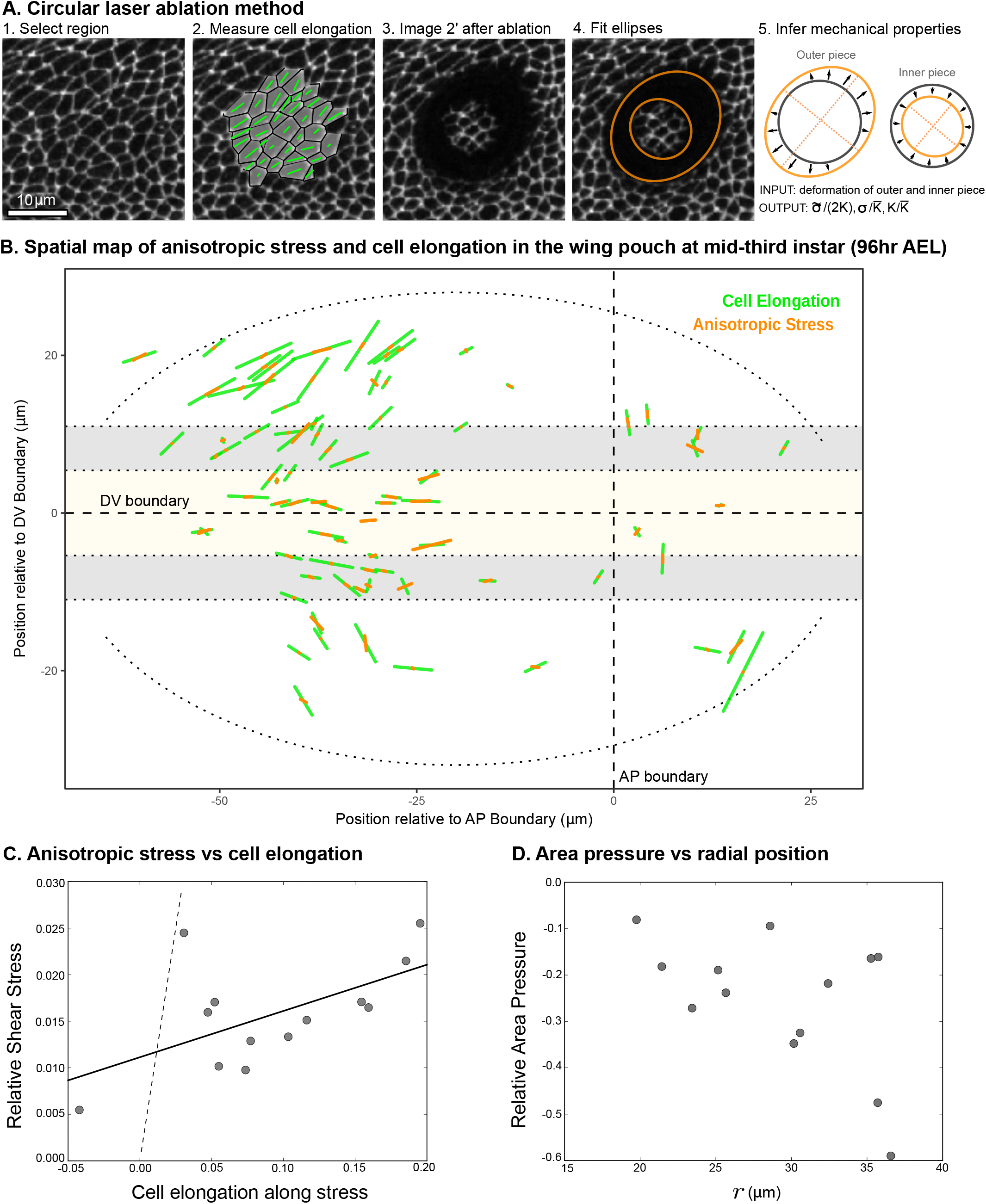
Circular laser ablation reveals patterns of tissue stress. (A) Description of the circular laser ablation method ESCA. The tissue is cut in a circle (radius 7*μm*), and cell elongation is averaged in this region before the cut. After the tissue relaxes (2 *min*), we fit ellipses to the cut region. We then infer mechanical properties using our model, which inputs deformation and outputs stress as a function of elastic constant and the ratio of the isotropic and anisotropic elastic constants (see also Fig S4). (B) Map of stress and cell elongation in the wing disc (96hr AEL). Each line represents a nematic, where length indicates the magnitude and angle indicates its orientation. Each data point comes from a different wing disc. The grey region indicates cuts that are straddling the border of two regions and were not analyzed further. The dotted lines on either side of the DV boundary indicate our cut-offs for delineating the border and DV boundary regions (see Methods). (C) Magnitude of anisotropic stress 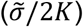 is plotted against cell elongation (projected onto the stress axis) for all ablations outside the DV boundary and border regions. Dotted line indicates a line with slope = 1, corresponding to a tissue lacking a nematic cell polarity cue. Fit line is solid black (See Theory Supplement part II). Data for the DV boundary region is presented in Fig S4. (D) Relative area pressure is plotted against *r* for all ablations outside the DV boundary and border regions. The correlation coefficient = −0.52.

The relationship between cell elongation and stress normalized by the elastic modulus has a slope 1 in the absence of polarity-driven stress (see Eq 2). We observe a much smaller slope for this relationship in our data (Fig 4C), indicating that polarity-driven stress is significant. We now use these data to estimate the parameters of our mechanical model. We write the shear stress defined in Eq 2 in terms of cell elongation and cell rearrangements, eliminating the orientational cue *q_rr_* using Eq 3. For the radial components, we have:

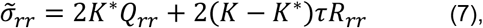

where *K*\* = (1 − *ζ*/(2*Kτλ*))*K* is an effective shear elastic coefficient. The difference between *K*\* and *K* depends on the parameters *ζ* and *λ* associated with the nematic cell polarity. We fit Eq 7 to the data and find *K*\*/*K* = 0.05 ± 0.02 and (1 − *K*\*/*K*)*τR_rr_* = 0.011 ± 0.002 (Fig 4C, see Theory Supplement part II). Combined with data from Fig 1, we find an estimate for the tissue relaxation time *τ* = 2 ± 2 fors, which is roughly consistent with that found during pupal morphogenesis (Etournay et al., 2015). From our data, we can also infer the radial profile of tissue area pressure, revealing that pressure increases towards the center (Fig 4D and Theory Supplement). This finding is consistent with the observed cell area profile, with smaller cell areas towards the center (Fig 1B,E).

In sum, we find a stress profile in the wing disc that is consistent with the observed measurements of both cell elongation and area. Further, we use these data to measure certain parameters of our biophysical model, including the tissue relaxation timescale and the effective shear elastic coefficient.

### Reduction of planar cell polarity pathways does not reduce tangential cell elongation

In our model, the radial orientation cue is required to generate the observed patterns of cell morphology, cell rearrangement, and tissue stress. Candidates for such an orientational cue are the planar cell polarity pathways (PCP), which are groups of interacting proteins that polarize within the plane of the epithelium. There are two well-characterized PCP pathways: Fat and Core (Butler & Wallingford, 2017; Eaton, 2003). In the wing, these systems form tissue-scale polarity patterns during growth (Brittle et al., 2012; Merkel et al., 2014; Sagner et al., 2012) and are required to position the hairs and cuticle ridges on the adult wing (Adler et al., 1998; Doyle et al., 2008; Eaton, 2003; Gubb & Garcia-Bellido, 1982; Hogan et al., 2011). To determine whether either of these pathways could function as the orientational cue described in our model, we analyzed cell elongation patterns after their removal.

We perturbed the Fat pathway using Nub-Gal4 to drive the expression of RNAi constructs targeting both Fat and Dachs (D) in the pouch region throughout the third larval instar. Consistent with previous work showing that loss of both Dachs and Fat suppresses wing growth (Cho & Irvine, 2004), we observe a significantly smaller wing pouch upon Fat/D double knockdown (Fig 5A). Nonetheless, the pattern of tangential cell elongation persists to the end of larval development (Fig 5A-C). Using scaled coordinates, we find that the radial profiles of cell elongation in Fat/D RNAi and control wings are similar (Fig 5D).

**Figure 5:**
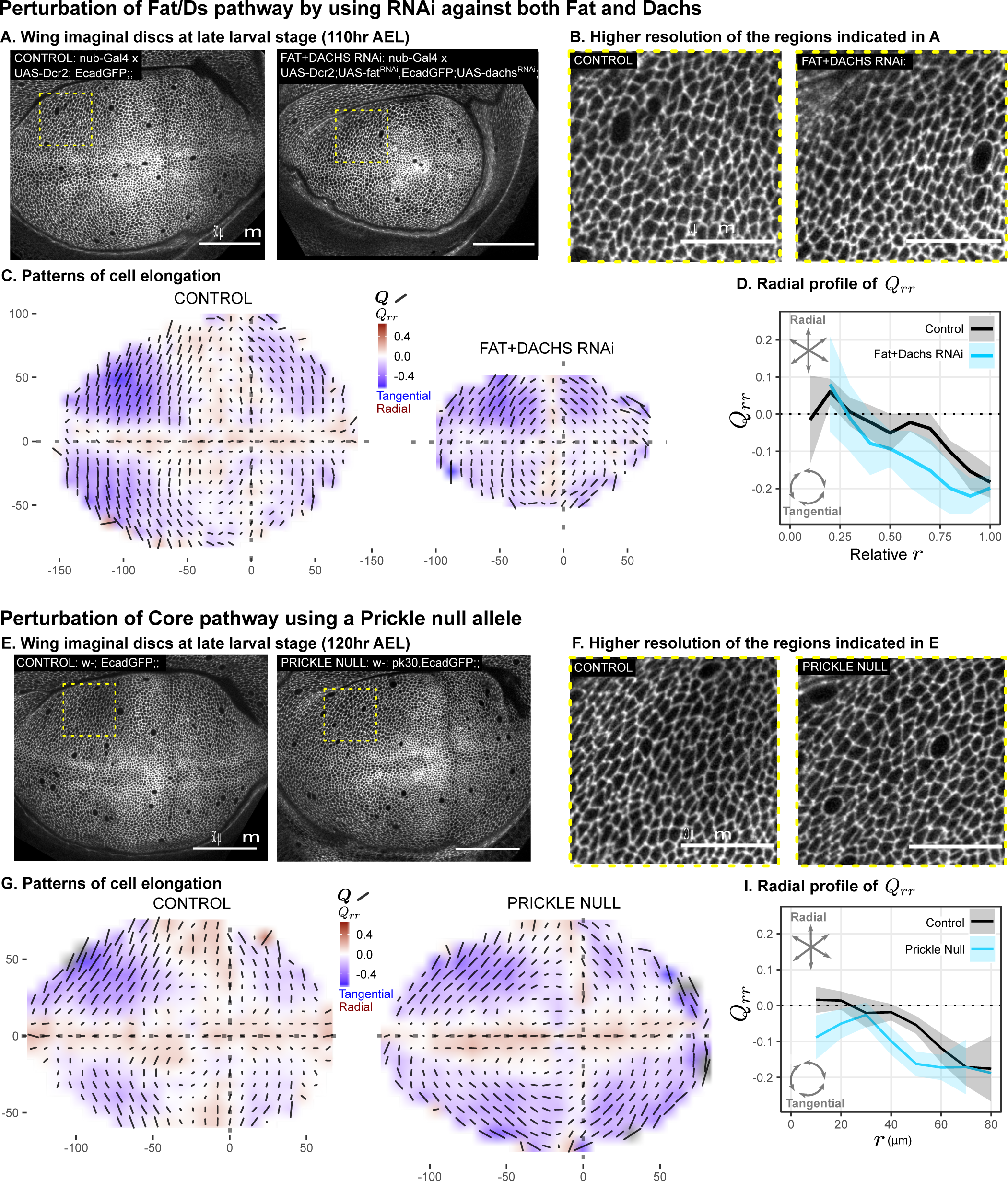
Reduction of planar cell polarity pathways does not reduce tangential cell elongation. (A-D) RNAi was induced against both Fat and Dachs in the pouch (nub^GAL4^ > UAS-Dcr, UAS-fat^RNAi^, UAS-dachs^RNAi^), and cell elongation patterns at the end of larval development (110hr after egg laying) are presented along with control (nubGAL4 > UAS-Dcr). (E-I) Cell elongation patterns at the end of larval development are presented for a null mutant in Prickle (*pk*^30^) and the wild type control. (A,E) Representative images of each genotype, with apical cell boundaries marked with Ecadherin-GFP. Yellow box indicates the inset that is presented in higher resolution in (B,F). (C,G) Cell elongation averaged across several discs for each genotype in a grid centered on the AP and DV boundaries. Bars represent the average cell elongation tensor ***Q***, where the length of the bar is proportional to the magnitude of cell elongation, and the angle indicates its orientation. Color indicates the radial component of cell elongation *Q_rr_*. Axis labels indicate distance from the AP (X) or DV (Y) compartment boundaries (in *μm*). (D,I) The radial profile of cell elongation for each genotype was quantified by averaging *Q*_rr_ in radial bins and plotting as a function of *r*. Since the fat/d RNAi wings are smaller, we present this profile as a relative distance to the center. The band of cells around the DV boundary was removed, and because the fat/d RNAi discs are smaller, this region of exclusion is a larger relative distance. Plots in (C-D) represent averages of *N* = 7 − 8 wing discs per genotype; for plots in (G-I), *N* = 11 − 14 per genotype.

We perturbed the Core PCP pathway using a previously characterized null mutation in Prickle (*pk*^30^), which causes defects in adult wing hair orientation (Gubb et al., 1999). We found that the cell elongation pattern in the Pk mutant is similar to the wild type control (Fig 5E-I). In the Pk mutant, the region of tangential cell elongation extends even further into the center than in control wings.

We conclude that the tangential cell elongation pattern persists in the absence of either PCP pathway. This result excludes these pathways as orienting cues for the cell elongation patterns.

### Mechanosensitive feedback generates self-organized patterns of cell morphology

We have shown that perturbing PCP pathways does not affect the radial patterns of morphology, raising the question of how orientational cues might arise. In previous sections, we have considered the orientational cue to be provided by a cell polarity system that is independently patterned. However, cell polarity in general would be affected by stresses in the tissue. Indeed, there are many examples of cells polarizing in response to mechanical stress (Duda et al., 2019; Hirashima & Adachi, 2019; Ladoux et al., 2016; Ohashi et al., 2017). Here, we show that introducing mechanosensitive feedback to the model of tissue mechanics can give rise to spontaneous emergence of the cell polarity cue (Fig. 6).

**Figure 6:**
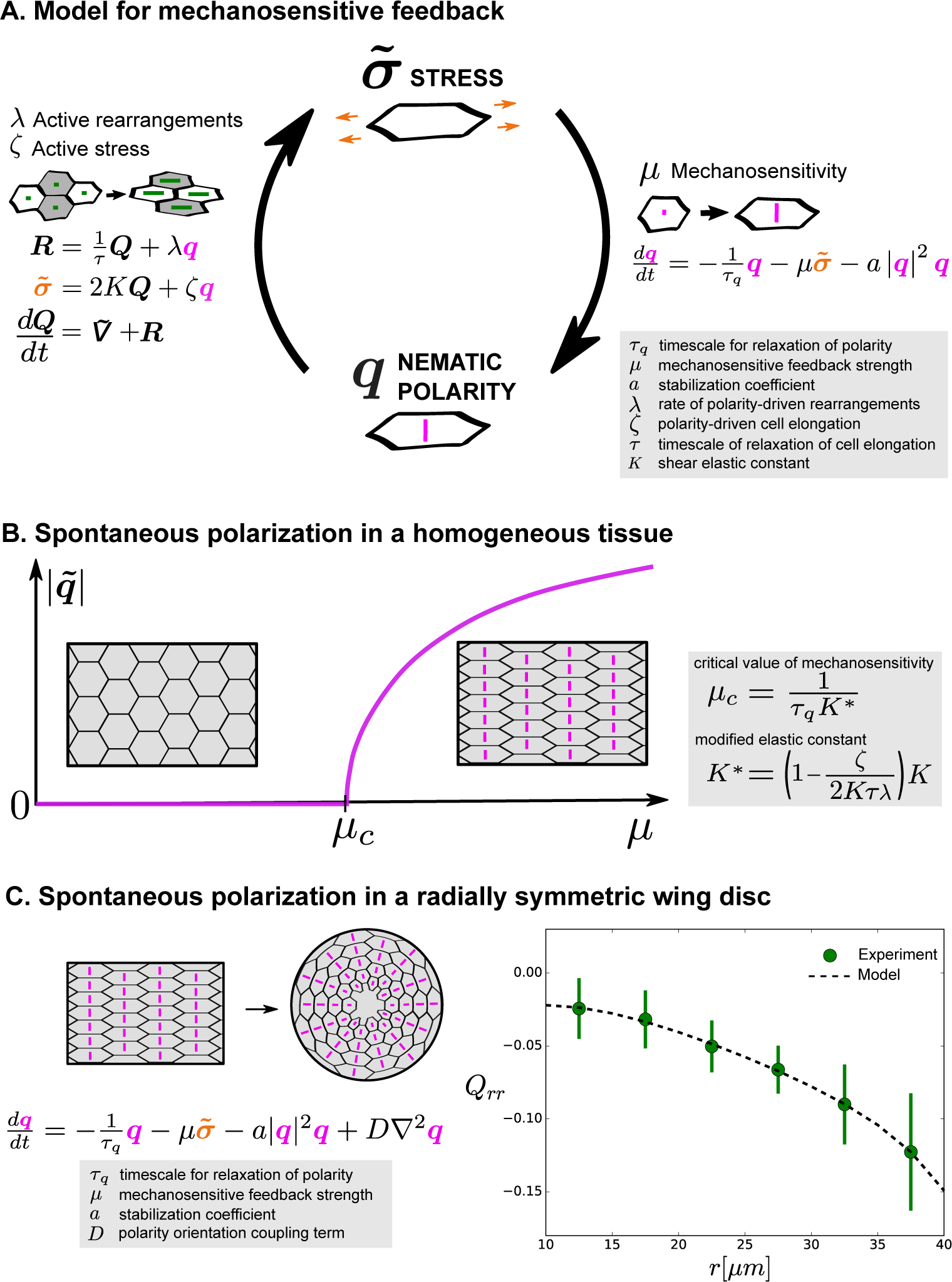
Mechanosensitive feedback generates self-organized patterns of cell morphology. (A) We introduce mechanosensitive feedback into our model with a dynamic equation relating the orientational cue ***q*** to tissue stress. With a change in ***q***, there can be a change in polarity-driven cell rearrangements and stress. (B) Above a critical value of *μ*, the isotropic state is unstable and the system spontaneously polarizes. (C) Application of the model to the radially symmetric wing disc. Generation of large-scale tissue polarity requires the orientation coupling term *D*. We find a set of parameters that can account for the experimental data on cell elongation in the wing disc (See Theory Supplement part ID and Table 1).

Mechanosensitivity is incorporated into our model of tissue mechanics through a dynamic equation for the orientational cue *q* that becomes stress-dependent:

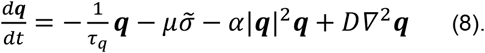

Here, ***τ**_q_* is a relaxation timescale for ***q**, μ* is a mechanosensitive feedback strength, the coefficient *α* > 0 ensures stability, and *D* is a coupling strength locally aligning orientational cues.

Now, Eq 8 provides a mechanosensitive feedback to Eqs 1, 2 and 3. These combined equations show a novel behavior. In particular, the orientational cue can emerge spontaneously by self-organization (Fig 6A-B). Beyond a critical value *μ_c_* of the mechanosensitive feedback strength *μ*, an isotropic tissue with *q* = 0 is no longer stable, and a state with an orientational cue *q* ≠ 0 emerges instead (Fig 6B and Theory Supplement part ID). The magnitude of this spontaneous polarization is |***q***| = *q*_0_, where 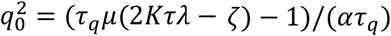, where a positive coefficient *α* is needed to stabilize the polarized state. By this mechanism, the anisotropic cue introduced earlier in our model can be locally generated by mechanosensitive self-organization and does not require the existence of pre-patterned polarity cues. To generate a large-scale pattern from locally generated anisotropic cues, they need to be aligned in neighboring regions. This local alignment is captured in Eq 8 by the orientation coupling term with strength *D*, which is similar to alignment terms found in anisotropic physical systems, such as liquid crystals (Gennes & Prost, 1993; Jülicher et al., 2018; Marchetti et al., 2013).

To discuss cell morphology profiles in the wing disc, we consider a simplified tissue model with radial symmetry, where the rate of radial cell rearrangement *R_rr_* is given (as estimated in Theory Supplement part IB) and the cell shape pattern and tissue stress pattern are calculated. Using a fit of cell elongation to the experimental data, we find a set of parameter values that accounts for the observed cell elongation patterns in the wing disc (Table 1, Theory Supplement). From this cell elongation pattern also follows the cell area pattern (as described above, Fig 3).

**Table 1:** Parameter values obtained by fitting the tangential cell elongation profile shown in Fig 6C to the self-organized model. Values in the last two rows are boundary conditions that are required to calculate the tangential cell elongation profile. The reported parameter uncertainty intervals were obtained by fitting 101 uniformly sampled tangential cell elongation profiles from the range defined by the standard deviation in the experimental tangential cell elongation profile (error bars in Fig 6C). The interval limits are the 10^th^ and 90^th^ percentiles of the obtained values.

| Fit parameter | Parameter value | Uncertainty interval |
| --- | --- | --- |
| $q_0^2$ | 0.01 | $[-0.06, 0.11]$ |
| $\lambda\tau$ | 1.0 | $[0.2, 1.3]$ |
| $\frac{D}{\alpha}$ | $0.5 \mu m^2$ | $[0.2, 27] \mu m^2$ |
| $\frac{1}{D} \left( \frac{1}{\tau\lambda} + 2K\mu\tau \right) R_{rr}$ | $9 \cdot 10^{-4} \mu m^{-2}$ | $[0.0, 3.2] \cdot 10^{-3} \mu m^{-2}$ |
| $q(r_{in})$ | 0.024 | $[0.014, 0.105]$ |
| $\partial_r q(r_{in})$ | $-1.2 \cdot 10^{-3} \mu m^{-1}$ | $[-9.3, 0.0] \cdot 10^{-3} \mu m^{-1}$ |

We conclude that our mechanosensitive model can account for the radial pattern of cell morphology in the wing disc. Note that because of the relatively large number of parameters used to fit a single experimental curve, there are large uncertainties when estimating parameter values. Nonetheless, the qualitative prediction that reduction in mechanosensitivity *μ* would lead to less polarization and thereby reduced cell elongation is rather robust and thus not likely to be affected by these uncertainties (see Theory Supplement part ID). We next test this prediction of our model experimentally.

### Suppression of mechanosensitivity weakens the gradients in cell elongation and cell size

In order to test our prediction that the reduction of mechanosensitivity will reduce the magnitude of cell elongation, we used RNAi to knockdown Myosin VI, a molecular motor implicated in mechanosignaling. Myosin VI is an upstream component of a Rho-dependent signaling pathway that reorganizes the actin-myosin cytoskeleton in response to mechanical stress (Acharya et al., 2018). Experiments in wing discs also indicate that mechanosensation involves Rho polarization and signaling (Duda et al., 2019). We performed RNAi against MyoVI in the wing pouch using Nub-Gal4 and evaluated cell morphology at the end of larval development (~120 *hr* AEL). We observe a clear reduction in the magnitude of tangential cell elongation as compared to wild type at this stage (Fig. 7A-D, Fig S5). In addition, our model predicts that such reduction of cell elongation would result in an increase of cell area in the central parts of the wing (Eqs 1-6). The observed pattern of increased cell area in MyoVI knockdown experiments is consistent with this prediction (Fig 7E-F). Therefore, the qualitative predictions of our model upon reducing the mechanosensitive feedback strength *μ* are confirmed by the experimental downregulation of the mechanosensitive motor, MyoVI.

**Figure 7:**
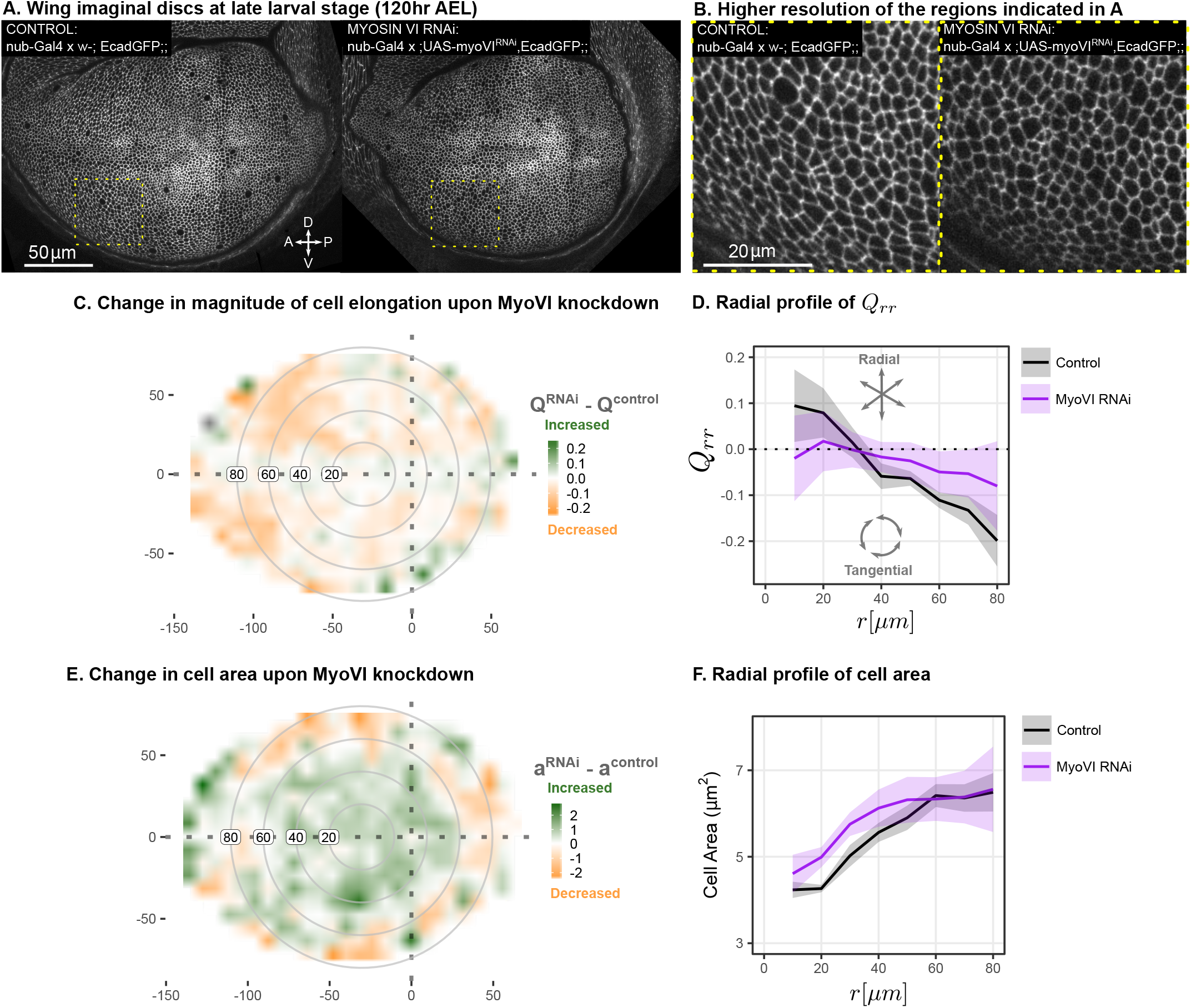
Suppression of mechanosensitivity weakens the gradients in cell elongation and cell size. (A) RNAi was induced against Myosin VI in the pouch (nub^GAL4^ > UAS-myoVI^RNAi^), and cell elongation and cell area were assessed at the end of larval development (119 *hr* after egg laying). Shown in (A) are representative images, with apical cell boundaries marked by Ecadherin-GFP. Yellow box indicates the inset that is presented in higher resolution in (B). The difference in magnitude of cell elongation (C) or cell area (E) between myoVI^RNAi^ and the corresponding control is presented. Axis labels indicate the distance to the AP boundary (X) or DV boundary (Y) in *μm*. Grey circles indicate radial bins, with numbers corresponding to distance (in *μm*) from the center. (D,F) The radial profile in the radial component of cell elongation *Q_rr_* (D) or cell area (F) was quantified for each genotype. Plots in (C-F) represent averages of *N* = 6 − 7 wing discs per genotype.

## DISCUSSION

Here, we have shown that patterns of cell shape and stress in the mid-third instar *Drosophila* wing disc do not rely on PCP pathways or differential growth. Instead, radially-oriented T1 transitions and tangential cell elongation emerge via mechanosensitive feedback in a self-organized process. We have presented a continuum model of tissue dynamics for this self-organization based on a mechanosensitive nematic cell polarity that accounts for the observed patterns of cell area, T1 transitions, and cell shape. Our work highlights a mechanism for the self-organized emergence of cellular patterns in morphogenesis.

### A pattern of T1 transitions is critical for cell morphology patterning in the *Drosophila* wing

Our work shows that the spatial pattern of T1 transitions is an integral part of the emergence of tissue organization during wing development. In contrast to situations such as germband extension, where T1 transitions exhibit clearly discernible patterns, the patterns of T1 transitions in the wing disc have been elusive. Many T1 transitions occur in the tissue in seemingly random orientations. However, on average, they exhibit a spatial pattern. We revealed these patterns by quantifying the nematics of T1 transitions and cell shape changes using the previously-described triangle method (Merkel et al., 2017) and then quantified them with radial averaging (Fig 2A-B). In this way, we revealed that a radial pattern of T1 transitions is linked to a tangential pattern of cell elongation.

Given this radial pattern of T1 cell rearrangements, the observed cell morphology pattern follows from a continuum tissue model based on a radially-oriented nematic cell polarity field (Fig 3). The polarity-oriented radial T1s create a cell shape pattern with corresponding patterns of tissue stress and tissue area pressure. The 2D area pressure is higher in the center and is lower towards the periphery. Note that this pressure profile does not rely on differential proliferation, as was previously proposed (Mao et al., 2013) but instead relies on a radial pattern of T1 transitions. We test our model using a novel circular laser ablation method. This method allows us to determine specific combinations of tissue parameters. In particular, we estimate the ratio of elastic constants 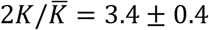 and *K*\*/*K* = 0.05 ± 0.02, as well as the cell shape relaxation timescale *τ* = 2 ± 2 *hr*.

This analysis raised the question of which nematic cell polarity cues guide the cell rearrangement and cell elongation patterns. PCP pathways are required for the proper orientation of T1 transitions in other contexts (Bosveld et al., 2012). However, we found that neither of the two known PCP pathways in the wing are required for the observed tangential cell elongation patterns (Fig 5). We instead show that an orientation cue can arise through self-organization via mechanosensitivity and identify MyoVI as a key molecular player.

### Mechanosensitive feedback can create self-organized patterns of cell morphology

Cell polarity cues can emerge via mechanosensitive feedback by transforming mechanical cues into chemical anisotropies. Nematic cell polarity can then orient active stresses and thereby amplify the mechanical stimulus (Fig 6). We introduce this mechanosensitive feedback in our continuum theory, which quantitatively describes the emergence of patterns of cell shape and cell rearrangements. The strength of this mechanosensitive feedback is described by a parameter *μ*. If *μ* exceeds a critical value *μ_c_*, an orientation cue and elongated cell shapes spontaneously emerge, while for *μ* < *μ_c_*, the tissue remains isotropic (Fig 6B). This model can account for the observed patterns of cell area and cell elongation in the wing disc and predicts that the reduction of mechanosensivity will result in reduced cell elongation. To test this prediction, we perturbed a RhoA-dependent mechanotransduction pathway by knocking down an upstream component, MyoVI, using RNAi (Fig 7). We find a clear phenotype of reduced cell elongation and increased cell areas in the center region, as predicted by our model.

This result, together with the fact that MyoVI is involved in Rho-dependent activation of actin-myosin cytoskeleton (Acharya et al., 2018), suggests that MyoVI is a molecular component of the mechanosensitive feedback we describe in our self-organized model. However, the molecular nature of the cue that defines the nematic cell polarity is unknown. This cue may organize the structure or dynamics of the actin-myosin cytoskeleton or the actin-myosin system itself could define nematic cell polarity. Indeed, it has been shown that myosin II localizes to long cell boundaries in the wing (LeGoff et al., 2013), corresponding to a nematic polarity aligned with the nematic cell polarity ***q***. Also, wing disc stretching experiments have shown that MyoII can polarize in response to exogenous stress in a Rho-dependent manner (Duda et al., 2019). Precisely how the actin-myosin cytoskeleton is affected by MyoVI in this system and how these cytoskeletal elements together guide cell rearrangements in response to anisotropic tissue stresses and cell shape changes remain open questions for future research.

Lastly, our laser ablation analysis shows that the region around the DV boundary has a different ratio of elastic constants than the rest of the tissue, which could affect the self-organized pattern formation we describe. Therefore, it will be interesting to study how Wingless/Notch signaling, which defines the DV boundary, may influence the mechanical properties that lead to mechanosensitive self-organization of polarity and morphology. In addition, we observe a richer pattern emerging very late in development (see Fig S5, Fig 5), including a region anterior to the AP boundary that is radially elongated. Future research will expand upon the model presented here to explore the dynamics of these patterns.

In summary, we used the *Drosophila* wing disc to identify a mechanism by which tissue morphology can arise from the self-organization of a mechanical feedback coupling cell polarity to active cell rearrangements. This mechanism is general and could be employed in other tissues and organisms to generate patterns of cell shape and cell area. Thus, we hope our work inspires new avenues of research that integrate theory and experiment to understand biological self-organization.

## ACKNOWLEDGEMENTS

We thank Stephan Grill for the microscope used to perform circular laser ablations and to Romina Piscitello-Gomez for assistance with these experiments. We thank the Light Microscopy Facility of the MPI-CBG for assistance with all other imaging experiments. Benoit Lombardot, formerly of the Image Analysis Facility of the MPI-CBG, provided us with a FIJI script to perform the projection of the apical surface, originally designed by Dagmar Kainmueller. Franz Gruber generated the *pk30,ecadGFP* fly line. We thank Christian Dahmann, Charlie Duclut, Kinneret Keren, Yonit Maroudas-Sacks, Alphee Michelot, Ioannis Nellas, and Romina Piscitello-Gomez and for critical review of the manuscript. This work was supported by funding from the Deutsche Forschungsgemeinschaft (EA4/10-1, EA4/10-2; N.A.D., K.V.I, S.E.), Swiss National Science Foundation (Grant #200021-165509; M.P.), and the Simons Foundation (Grant #454953 to Matthieu Wyart; M.P.). Lastly, we dedicate this manuscript to our co-author, Suzanne Eaton, who passed away tragically towards the conclusion of this work.

## AUTHOR CONTRIBUTIONS

Conceptualization: N.A.D., M.P., S.E., F.J.; Methodology: N.A.D., K.V.I, M.P., S.E., F.J.; Software: N.A.D., M.P.; Validation: N.A.D, M.P.; Formal Analysis: M.P.; Investigation: N.A.D., K.V.I; Resources: N.A.D.; Data Curation: N.A.D.; Writing – Original Draft: N.A.D., M.P., F.J.; Writing – Review and Editing: N.A.D., M.P., K.V.I., F.J.; Visualization: N.A.D., M.P.; Supervision: S.E., F.J.; Project Administration: N.A.D., M.P., S.E., F.J.; Funding Acquisition: S.E., F.J.

## MATERIALS AND METHODS

### LEAD CONTACT AND MATERIALS AVAILABILITY

This study did not generate any new unique reagents. All requests for further information and reagents may be directed to the lead author.

### EXPERIMENTAL MODEL AND SUBJECT DETAILS

All experiments were performed with *Drosophila melanogaster*, using lines that are publicly available and previously published. Our *Drosophila* lines were fed with a standard media containing cornmeal, molasses agar and yeast extract and grown under a 12 h light/dark cycle. All experiments were performed at 25 °C. Both males and females were analyzed, and the sex of the animals was not recorded, as we have no reason to believe there is any sexual dimorphism in the studied phenomenon. To synchronize development, we collected eggs deposited within a defined time window on apple juice agar plates. To do so, we transferred the flies from standard food vials to cages covered by apple juice agar plates containing a dollop of yeast paste for food. After at least 2 *hrs*, the plates were replaced, and the timing of collection started. Eggs laid within a ~2 *hr* time window were collected by cutting out a piece of the agar and transferring it to a standard food vial. We limited the number of eggs per vial to < 15 to avoid crowding. The middle of the time window for egg collection was considered to be 0 *hr* after egg laying (AEL). Experiments from timelapse imaging and laser ablation (Figs 1, 2, and 4) were captured after explanting at 96 *hr* AEL, whereas those involving RNAi (Figs 5 and 7) were explanted at 110 – 120 *hr* AEL to allow the maximal amount of time for the RNAi phenotype to emerge. The specific genotypes used for each experiment are indicated in the figure legends and in the Key Resources Table.

### METHOD DETAILS

#### Timelapse image acquisition and processing

##### Sample preparation

Wing explants were grown *ex vivo* as described previously (Dye et al., 2017). Briefly, wing discs were dissected from larvae in growth media (Grace’s cell culture media + 5% fetal bovine serum + 20nM 20-hydroxyecdysone + Penicillin-Streptomycin) at room temperature. Then, they were transferred to a Mattek #1 glass bottom petri-dish to the center of a hole cut in a double-sided tape spacer (Tesa 5338) and covered with a porous filter. The dish was then filled with fresh growth media.

##### Acquisition

Data from movies 1-3 were used previously (Dye et al., 2017). Movie 4 was acquired after publication of the first manuscript, in the same exact way as movies 1-3. Briefly, E-cadherin-GFP-expressing wing discs were imaged in growth media using a Zeiss spinning-disc microscope to acquire 0.5 *μm* spaced Z-stacks at 5 *min* intervals with a Zeiss C-Apochromat 63X/1.2NA water immersion objective and 2×2 tiling (10% overlap). This microscope consisted of an AxioObserver inverted stand, motorized xyz stage, stage-top incubator with temperature control set to 25C, a Yokogawa CSU-X1 scanhead, a Zeiss AxioCam MRm Monochrome CCD camera (set with 2×2 binning), and 488 laser illumination. We circulated growth media during imaging using a PHD Ultra pump (Harvard Apparatus) at a rate of 0.03 *ml*/*min*. Time 0 *hr* of the movie was considered to be the start of imaging acquisition, typically 45 – 60 *min* from the start of dissection (time required for sample and microscope preparation).

Movie 5 was acquired on a newer microscope with a larger field of view that eliminated the need to tile across the wing pouch. This microscope has an Andor IX 83 inverted stand, motorized xyz stage with a Prior ProScan III NanoScanZ z-focus device, a Yokogawa CSU-W1 with Borealis upgrade, and a Pecon cage incubator for temperature control at 25 °C. We used an Olympus 60x/1.3NA UPlanSApo Silicone-immersion objective with 488 laser illumination and an Andor iXon Ultra 888 Monochrome EMCCD camera. We acquired 0.5 *μm* spaced Z-stacks of a single tile at 5 *min* intervals, as for the other movies. Movie 5 was also acquired without the constant flow of new media.

For all movies, care was taken to limit light exposure, using laser power values of < 0.08 *mW* and exposure times less than 350 *ms* per image.

##### Processing

Raw Z-stacks were denoised using a frequency bandpass filter and background subtraction tools available in FIJI (Schindelin et al., 2012). Then, we used a custom algorithm as described previously (Dye et al., 2017) to make 2D projections of the apical surface, marked by Ecadherin-GFP. This algorithm also outputs a height-map image, in which the value for each pixel corresponds to the level in the Z-stack of the identified apical surface. For those movies that were tiled, we used the Grid/Collection Stitching FIJI plugin (Preibisch et al., 2009) to stitch the 2D projections, and then used that calculated transformation to stitch the height map images, so that we could correct the cell area and elongation values for local curvature (see below). We focus in this work exclusively on the disc proper layer, which is the proliferating layer of the disc that goes on to produce the adult wing. We did not consider cell shape patterns in the peripodial layer, the non-proliferating layer of the wing disc that is largely destroyed at the onset of pupariation.

#### Single timepoint image acquisition and processing for analysis of RNAi/mutant phenotypes

##### Sample preparation

Samples for single timepoint imaging were acquired exactly as described for live imaging: dissected in growth media and imaged in Mattek dishes under a porous filter.

##### Acquisition

All imaging of single timepoint data was performed with the same microscope described above for Movie 5. We did not acquire timelapse data for these genetic perturbations, and thus we chose to image the entire disc using 2×2 tiling, 0.5 *μm* spaced Z-stacks. Approximately 6-10 discs were imaged per dish, within approximately 30 – 60 *min*.

##### Processing

Tiled images were stitched together as Z-stacks; then we obtained the apical surface projection and its corresponding height map as described above and in (Dye et al., 2017).

#### Circular Laser ablation

##### Acquisition

Wing discs from mid-third-instar larvae (96 *hr* AEL) were dissected and mounted exactly as was done for the live imaging timelapses. Due to constraints on the speed of ablation, we only cut regions of the wing disc pouch that were completely flat, so that we could cut in a single plane at the apical surface (rather than having to cut in each plane of a Z stack). Where this flat region lies depends on how the disc happens to fall on the coverslip during mounting. Because the anterior compartment is larger and higher, most of our ablations are in this compartment. To access other regions, we also mounted some wing discs on an agarose shelf: stripes of 1% agarose (in water) were dried onto the surface of the coverslip and wing discs were arranged their anterior half propped up on the agarose shelf prior to adding the porous filter cover.

Ablations were performed using ultraviolet laser microdissection as described in (Grill et al., 2001) using a Zeiss 63X water objective. First, we took a full Z-stack of the sample prior to the cut. Then, we selected a 7 *μm* radius circle that would ablate in the flattest region of the tissue. No imaging is possible during ablation, but we acquired a 2 *min* timelapse immediately after the cut in a single Z-plane. This timelapse data was not used except to estimate whether or not the sample was fully cut. After, once the sample has finished expanding but not started to heal (~2 *min*), we took another full Z-stack to image the endpoint. We excluded a small number of data points if any of the following were true: (1) the inner piece remaining after the ablation was no longer visible (sometimes it floats or is destroyed); (2) the cut appeared to expand highly asymmetrically (rare); (3) the wing discs were clearly too young to be considered 96 *hr* AEL (poor staging).

##### Immunofluorescence after ablation

Due to a limited field of view on the microscope used for ablation, we performed immunofluorescence after the ablation in order to better estimate the position of the cut in the wing pouch. After all the discs in the dish were ablated (<10 discs/dish), the entire dish was fixed through the filter by adding 4% PFA and incubating 20min at room temperature. After, the dish was rinsed and kept in PBST (PBS + 0.5% Triton X-100) until all discs from that image acquisition day were completed (2 – 4 *hr*). All samples were then blocked using 1% BSA in PBST+250mM NaCl for 45 *min* – 1 *hr*, and then incubated in primary antibody overnight (diluted in BBX: 1% BSA in PBST). Initially, we labeled samples with SRF (Active Motif, 1:100 dilution), but later we switched to Patched and Wingless (DSHB, 1:100) to identify the compartment boundaries. Primary antibody toward GFP (recognizing the E-cadherin-GFP) was also included. After overnight incubation at 4C, we washed with BBX, followed by BBX+ 4% normal goat serum (NGS), for at least 1 *hr*. Secondary antibodies were added for 2 *hr* at RT in BBX+NGS. Finally, samples were washed 4×10min in PBST and imaged in this media. Imaging was performed on one of the two spinning disc microscopes described above for live imaging. We matched the stained samples with the ablation images by (1) keeping track of the position of each disc on the dish and where it was ablated during acquisition, and (2) morphology of the disc before/after ablation. All antibody information is listed in the Key Resources Table.

### QUANTIFICATION AND STATISTICAL ANALYSIS

#### Cell segmentation, tracking, and alignment

##### Segmentation and tracking

Using the 2D projections, we performed cell segmentation and tracking using the FIJI plugin, TissueAnalyzer (Aigouy et al., 2016). We manually corrected errors in the automated segmentation and tracking as much as possible and then generated a relational database using TissueMiner (Etournay et al., 2016). Images were rotated to a common orientation (Anterior left, dorsal up). We then manually identified three regions of interest at the last point in the timelapse, using the FIJI macros included with TissueMiner: the “blade” was defined roughly as an elliptical region surrounded by the most distal folds; and the Anterior-Posterior and Dorsal-Ventral boundaries were estimated using the brightness of E-cadherin-GFP and apical cell size (Jaiswal et al., 2006).

For the timelapse data, we used only the cells within these manually defined regions that were trackable during the entire course of the movies. An example of this region is presented for Movie 1 in Fig 1A. Furthermore, we also excluded from all of our analysis the first 2 *hr* of imaging, the so-called adaption phase, where cells uniformly shrink in response to culture (Dye et al., 2017).

For the RNAi data (Fig 5 and 7), we did not acquire timelapses, rather a single timepoint at late stages of development. We nevertheless chose to analyze these data using the same TissueMiner workflow for simplification. Because TissueMiner was developed for timelapse data, however, it requires at least two timepoints. Thus, we duplicated the data and labelled the two images as if they were timepoints 0 and 1 of a timelapse. We then only analyzed timepoint 0. The regions of interest in these data, therefore, are manually defined and not simply the region that is trackable (since it is static data).

##### Alignment on compartment boundaries

To average across all movies of timelapse data, or all discs of the same genotype of the static RNAi data, we generated a disc-coordinate system by normalizing the position of each cell to the AP and DV boundaries for that disc. To do so, we averaged the absolute xy positions of all the cells in the AP or DV boundaries over all time after the adaption phase. We then calculated the distance of each cell to the new X axis (average position of the AP boundary) and Y axis (average position of the DV boundary).

#### Analysis of cell size and shape

##### Definition of the cell elongation tensor

The cell elongation tensor ***Q*** is a traceless symmetric tensor that quantifies the anisotropy of cell shapes in a region of the tissue. We define cell elongation using a triangulation of the tissue obtained by connecting centroids of connected cells (Etournay et al., 2015; Merkel et al., 2017). The state of each triangle is described by a tensor ***s***, defined by a mapping an equilateral reference triangle to the triangles in the tissue. The state tensor contains information about the triangle elongation tensor 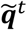, orientation angle *θ*, and area a:

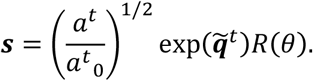

Here, 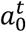 is the area of the reference triangle, and *R* is 2-dimensional rotation matrix. Cell elongation in the tissue region is defined as an area weighted average of triangle elongation. For details about the method see (Merkel et al., 2017).

##### Adjustment of cell area and elongation to account for tissue curvature

The wing disc pouch has a slightly domed shape (Fig S1A). After projection onto a 2D surface, the cell shapes and areas will be distorted. To ensure that the radial profiles we measure in a 2D projection (Fig. 1B,C,E, and F and Fig. 3F,G) are not a result of tissue curvature, we account for the distortion caused by projection. We use the height maps generated by our projection algorithm, which identifies the apical surface of the pouch within the 3D Z-stacks. We smooth this height map with a gaussian filter of width *σ* = 2 *μm* to find the height field *h*(*x,y*), and then we calculate the height gradient field. We then smooth the result again with the same gaussian filter to find 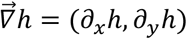. The deprojected cell or triangle area *a*_0_ is obtained from the measured area *a* as 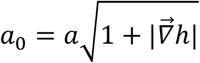, where 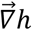 is evaluated at the cell center. To find the deprojected cell elongation of a triangle we evaluate 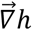 at the triangle center. Then, we find the angle of steepest ascent *α* = arctan(*∂_y_h*/*∂_x_h*) in the projected plane and define the tilt transformation:

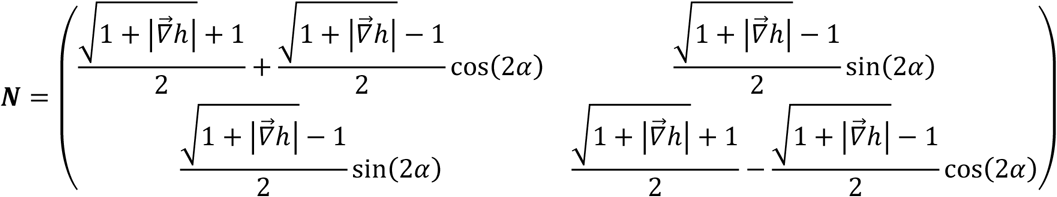

We apply this transformation to the triangle shape tensor ***s***, as defined in (Merkel et al., 2017), to determine the deprojected triangle shape tensor ***s***_0_

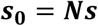

The cell elongation tensor ***Q*** is obtained from the triangle shape tensor ***s*_0_** as the corresponding area weighted average using the deprojected cell area, as described in (Merkel et al., 2017).

##### Spatial maps of cell size and shape

To generate the color-coded smoothed plots of area and elongation (Fig 1B-C), we divided the aligned wings into a grid with boxsize = 10pixels (~2 *μm*). For each position, we averaged cell area or performed an area-weighted averaged of triangle elongation in a neighborhood box = 20 pixels (~4 *μm*). To create the nematic elongation pattern, we similarly averaged elongation of triangles whose centers lie within each grid box, with grid box size = 30 pixels (~6 *μm*).

##### Calculation of radial elongation center point

We define the center of the wing pouch to be on the DV boundary. To determine the location of the center along the DV boundary *x_c_* we divide the pouch into four regions defined by the DV boundary and a line perpendicular to it located at some position *x*, as shown in Fig S1E. Then, we define a function

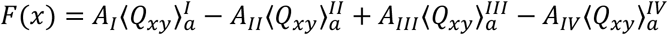

for all values of *x* along the DV boundary. Here, *A^I−IV^* are the areas of the four regions and 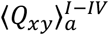 are the area weighted averages of the cell elongation component *Q_xy_* calculated in the four regions. Finally, *x_c_* is the value that minimizes *F*(*x*) (Fig S1F). We find that the center point lies just anterior to the intersection with the anterior-posterior (AP) boundary (Fig 1, S1, consistent with (LeGoff et al., 2013)).

In time-lapse experiments, cell centers were determined using the last 20 timepoints. In single timepoint image experiments (Figs 5 and 7), all images of a single genotype were used together after alignment on the DV and AP boundaries.

##### Exclusion of the band of cells near the DV boundary

In contrast to the other regions (blade, AP and DV boundaries), the definition of the band around the DV boundary (Fig S2) and the region of the blade that excludes this region (Fig 1D) were not defined manually using the FIJI scripts of TissueMiner. Rather, we defined them after the TissueMiner databases were generated, using the position of cells relative to the DV boundary in the last frame. Using the plots of the radial elongation on each disc, we estimated the width of the stripe region, and then filtered for cells included in and excluded from this region. Once these cells in the last frame were identified, we used the UserRoiTracking.R of TissueMiner to backtrack these two regions, producing a list of all cells belonging to this lineage that are traceable forward and backward in the movie.

For the static RNAi data, we used the average width of the band around the DV boundary from timelapse data as an estimate and excluded this region from the static images to analyze the radial gradients in cell area and elongation (Fig 5 and 7).

##### Plotting area and elongation versus distance to the center of elongation

For the timelapse data, we defined the radial elongation center (described above) for each movie and then calculated the distance away from this center for each cell. After excluding the band of cells around the DV boundary, we binned cells by radius by rounding to the nearest 10 *μm*. We also binned the movie across five equal time windows (~2 *hr*), excluding the adaption phase. Average cell area and area-weighted average of cell elongation were calculated for each of these bins in each time group for each movie. Including data from all 5 movies, we report the global average and its standard deviation (Fig 1E-F). For Fig 3, we averaged over the last ~5 *hr*. For the band of cells around the DV boundary (Fig S2), we binned cells in *x*, rather than in radius, and report the gradient along the *x* axis, defined as the DV boundary.

The static RNAi data were analyzed similarly. We report the global average of all discs in the genotype and the standard deviation (Figs 5 and 7). The number of discs analyzed in each genotype is listed in the figure legend.

#### Regional analysis of isotropic tissue deformation

##### Spatial maps of cellular contributions to isotropic tissue deformation

To analyze the pattern of isotropic deformation, we locally averaged cell behavior by dividing the tissue into a grid centered upon the crossing of the AP and DV boundaries. First, we determined the average position of cells in the AP and DV boundaries to generate a common frame of reference across all movies. Second, we divided the tissue visible in the last frame into a grid centered on these compartment boundary positions, with grid size = 8 *μm*. Grid boxes that were incompletely filled (less than 33% of the area of the box filled by cells) were discarded to eliminate noise along the tissue border. Third, each grid box was considered an “ROI” and then tracked through the entire movie using TissueMiner’s ROI tracking code. Fourth, the rate of deformation by each type of cellular contribution was calculated as a moving average (kernal = 11) for each grid position for each timestep. Last, we averaged these rates over all time points post-adaption period and plotted in space (Fig 1H,I,K,L).

##### Plotting the radial profile

To show tissue deformation as a function of distance to the center, we first calculated the distance to the center of symmetry for each cell. Second, we divided the tissue visible in the last frame into radial bins, rather than a grid, by rounding the radial position of each cell to its nearest 10 *μm*. Third, as we did for the grid, we defined each radial bin to be an “ROI” and then tracked the region forward and backward using TissueMiner’s ROI tracking code. Fourth, the rate of deformation by each type of cellular contribution was calculated as a moving average (kernal = 11) for each radial bin ROI at each timestep. Last, we averaged these rates over all time points post-adaption period and plotted this rate as a function of distance to the center. Note that we also show the spatial profiles of tissue growth in the band around the DV boundary in Fig S2, but here we binned along *x*, not *r*.

#### Regional analysis of anisotropic tissue deformation

##### Spatial maps of cellular contributions to anisotropic tissue deformation

We previously published patterns of cellular contributions to anisotropic tissue shear from movies 1-3 (Dye et al., 2017); however here, we calculated these patterns in a more accurate way and report averages over multiple movies (Fig 2A). Previously, we assigned a grid at each timepoint as in (Etournay et al., 2015, 2016). While this method provides a simple first approximation of the pattern, it is not completely accurate because we are not tracking the box in time but reassigning it at each timepoint; thus, cell movement in/out of the box is not counted. Here, we perform grid box tracking, as described above for the calculation of cellular contributions to isotropic tissue deformation but with grid box size = 15 *μm*. Further, we present a global average of not just movies 1-3, but also the two new movies. We calculated for each movie the rate of tissue deformation by each cellular contribution as a moving average (kernal = 11) in time. Then, we averaged across all five movies at each timepoint and presented the pattern as a cumulative sum, starting from the end of the adaption phase (first 2 *hr*) and continuing through the end of the timelapse. All correlation terms were added together (see (Merkel et al., 2017)). The contribution to tissue shear from cell extrusion is very small and thus not shown.

##### Accumulating cellular contributions to radial tissue shear

We calculated a moving average (kernal size = 11) total value for each cellular contribution to radial tissue shear, averaged across the blade (excluding the band around the DV boundary). Then, we accumulated the contributions after the adaption phase (first 2 *hr*) and plotted over time (Fig 2B). In addition to the previously described types of correlations (Merkel et al. 2017), the radial decomposition also involves a correlation between cell area and shear (see Theory Supplement part III). We added all correlation terms together in Fig 2B.

#### Quantification of Circular Laser Ablation

##### Measurement of Stress, Cell Elongation, and Cell Area

For the ablation data, we first projected the Z-stack images of the wing discs before and after ablation using our custom surface projection (Dye et al., 2017).

To quantify cell elongation and area, we used the images taken before the cut. Cells were segmented using the FIJI plugin TissueAnalyzer and then processed with TissueMiner to rotate the disc to a common orientation (anterior to the left; dorsal up) and to create a triangle network and a database structure. The area-weighted average of triangle elongation and the average cell area was calculated for those cells included in the center of a circle of a radius = 1.3*cut radius (4.55 *μm*) centered at the center of the cut. Varying the size of this region only slightly affects the result: <1.0*cut radius, the distribution becomes more noisy because we are averaging a smaller region with less cells, but regions that are too big (~1.8*cut radius) may begin to include cells that span different regions of the wing (i.e., the band around the DV boundary). In Fig 4C and Fig S4B,E, we plot cell elongation projected onto the stress axis.

To measure the shape of the tissue after ablation, we fit ellipses to the inner and outer piece left by the cut using the projected after-cut images. These images were first rotated using the transformation performed on the before-cut images to orient the tissue. We used Ilastik (v. 1.2.2) (Berg et al., 2019) to segment the cut region: we trained the classifier using all the data from a single day’s acquisition, delineating three regions: cells, membranes, and dark regions. Using the trained classifier, we then generated a thresholded binary image of the cut tissue’s shape and cropped the image around the cut. We then fit inner and outer ellipses to these images.

##### Grouping ablations into regions of the wing pouch

To quantify the position of the cut in the wing pouch, the immunofluorescence post-cut images were projected using maximum intensity. In FIJI, we manually measured (in *μm*) the distance from the center of the cut to the DV boundary and AP boundary. We noted that fixation causes the tissue to shrink. We estimated the extent of shrinkage using a subset of the samples in which the compartment boundaries were visible in the Z-stack taken of the live sample before the cut. We used FIJI to measure distances from the center of the cut to the compartment boundaries in both the pre-fix live images and in the post-fix immunofluorescence images for this subset. We measure a discrepancy indicating that the tissue shrinks by ~15%. Thus, we compensate for this shrinkage when measuring distances to the compartment boundaries in our analysis. We also noted the compartment (dorsal/ventral/anterior/posterior) in post-fix images and added a sign to the distances to the boundaries (ventral and posterior getting negative “distances”) to create the spatial map shown in Fig 4B.

To classify the cuts by position, we first estimated the width of the band around the DV boundary from the timelapse data to be ~22 *μm* (centered on the DV boundary). We then classified a cut as in the DV boundary if its cut boundary extended no more than 1.4 *μm* (20% of the 7 *μm* radius) outside this 22 *μm* horizontal stripe region. Likewise, cuts were classified as outside this region if it did not extend more than 1.4 *μm* into the 22 *μm* horizontal stripe. We excluded all cuts that appear to straddle the border between these regions (grey in Fig 4B). We also excluded those cuts with centers lying within 7 *μm* of the AP boundary, in case material properties at this boundary region are also different.

#### ESCA: Determination of stress from response to circular ablation

To obtain the normalized shear stress 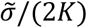, normalized isotropic stress 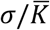 and the ratio of elastic constant 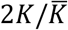, we fit ellipses to the inner and outer tissue outlines simultaneously. These three parameters determine the large and small semi-axes of the two ellipses. Other fitting parameters are the center positions of the two ellipses and the angles of major axes.

Fits were performed on all of the laser ablations in the same region (either within or outside of the band around the DV boundary) for a range of fixed values of the ratio 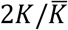. The optimal value was considered to be the one that minimizes the sum of fit residuals (Fig S4C,F). We defined a threshold of fit residual, as shown in Fig S4A, to eliminate cuts with non-elliptical outlines. The uncertainty of 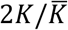 was estimated by bootstraping a subset of 7 cuts in each region 100 times; we report the standard deviation of the results.

### KEY RESOURCES TABLE

| REAGENT or RESOURCE | SOURCE | IDENTIFIER |
| --- | --- | --- |
| **Antibodies** |  |  |
| anti-*Drosophila* SRF mouse mAb, lot# 03504001 | Active Motif | Cat#39093 |
| anti-GFP rabbit IgG | Invitrogen | Cat#A11122 |
| anti-*Drosophila* Patched (mouse, Apa 1, conc) | Developmental Studies Hybridoma Bank | AB_528441 |
| anti-*Drosophila* Wingless (mouse, 4D4, conc) | Developmental Studies Hybridoma Bank | AB_528512 |
| **Chemicals, Peptides, and Recombinant Proteins** |  |  |
| Grace's insect medium | Sigma Aldrich | Cat#G9771-10X1L |
| Fetal Bovine Serum | Thermo-Fisher | Cat#10270098 |
| 20-hydroxyecdysone | Sigma Aldrich | H5142 |
| Penicillin-Streptomycin | Sigma Aldrich | P4333 |
| **Experimental Models: Organisms/Strains** |  |  |
| *Drosophila*: EcadherinGFP: yw-;EcadherinGFP;; | Suzanne Eaton lab MPI-CBG (Huang et al., 2009) |  |
| *Drosophila*: nub<sup>GAL4</sup>: w-;nubbin<sup>GAL4</sup>, EcadGFP;; | Suzanne Eaton lab MPI-CBG |  |
| *Drosophila*: fat+dachs<sup>RNAi</sup>:Dcr2;UAS-fatRNAi, EcadherinGFP;UAS-dachsRNAi | Vienna Drosophila Resource Center | Cat#9396, 12555 |
| *Drosophila*: pk30: w-;pk30,EcadherinGFP;; | Bloomington *Drosophila* Stock Center | Cat#44229 |
| *Drosophila*: myoVI<sup>RNAi</sup>: w-;UAS-myoVI<sup>RNAi</sup>,EcadGFP;; | Vienna Drosophila Resource Center | Cat#37534 |
| **Software and Algorithms** |  |  |
| TissueMiner | (Etournay et al., 2016) |  |

