## Supplement for "Self-organized patterning of cell morphology via mechanosensitive feedback"

### Fig S1, related to Fig 1: Quantification of the radial elongation pattern

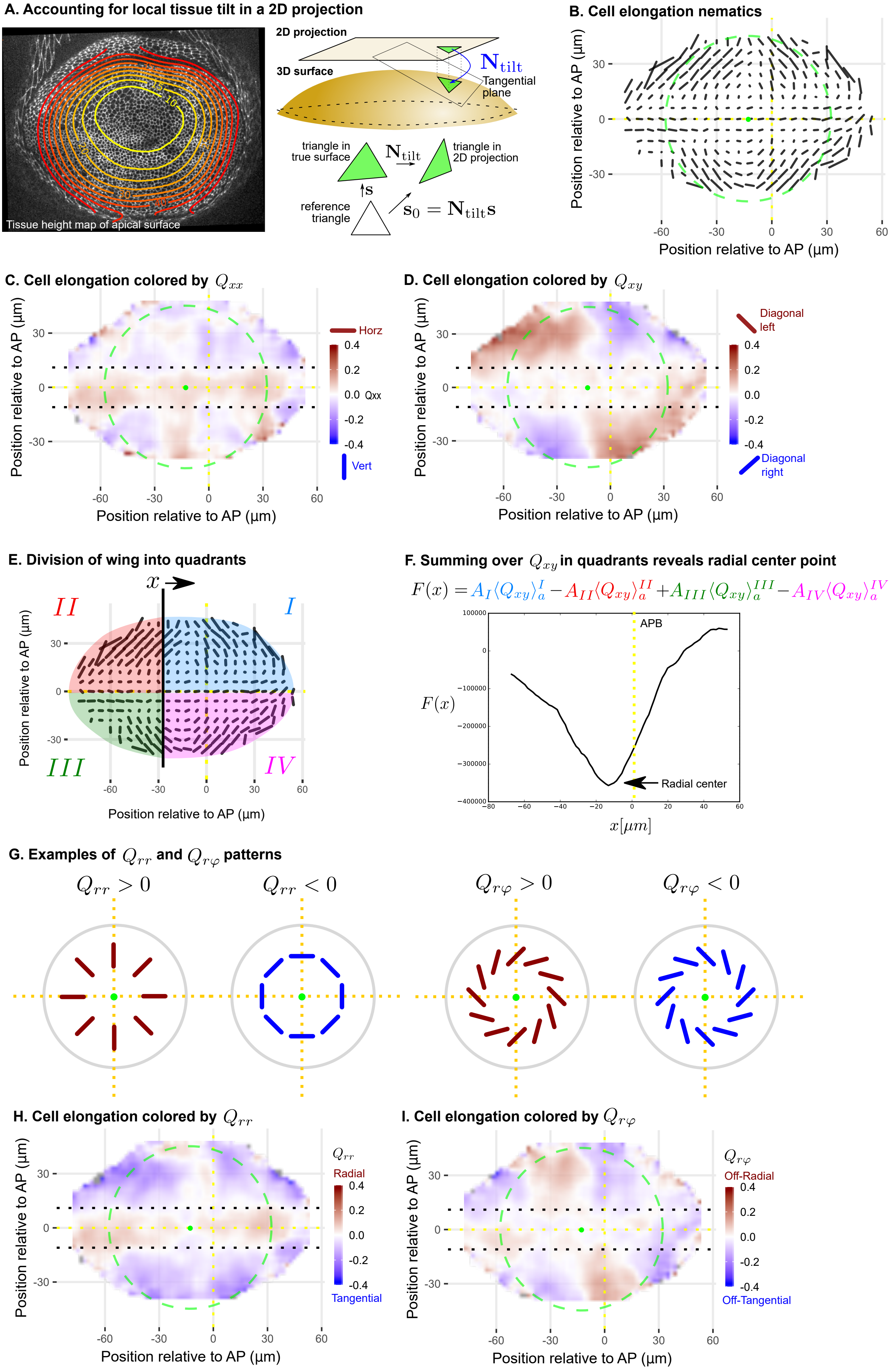

Analysis of cell elongation during timelapse of mid-third instar wing discs. (A) Cell area and elongation were adjusted to account for the distortion associated with projecting a curved surface in 2D. Left shows a 2D projected image from the timelapse movie of an E-cadherinGFP-expressing wing explant, overlaid with contours representing the depth of the apical surface in the acquired Z-stack. Numbers indicate distance (in um) from the most apical plane in the Z-stack. The right diagrams depict how we adjust the size and shape of the triange due to local tissue tilt. We determine the local tangential plane of the tissue surface at the triangle center. We then calculate the transformation matrix $N_{tilt}$ relating the shape tensor $s_0$ (in the tangential plane) and the observed shape tensor **s** (in the projected plane). (B) Cell elongation pattern presented in Fig 1, shown here without the color code. Cell elongation was averaged across all five movies in the middle time window within a grid centered on the AP and DV axes. The cell elongation nematic tensor **Q** is represented by bars, whose length is proportional to the magnitude and the angle indicates its orientation. The green circle is added to show the almost circular symmetry of the pattern, and the green dot is the calculated center of symmetry of the cell elongation pattern. (C-D) The cell elongation pattern shown in B, colored by the XX (B) or XY (C) component. (E-F) Calculating the center of symmetry of the cell elongation pattern. At each point along the x axis (DV boundary), we calculate the area-weighted sum of $Q_{xy}$ in the four quadrants. The center point lies at the function minimum, which lies just anterior to the AP boundary (APB). (G) Illustrations of the orientation of cell elongation for positive and negative values of $Q_{rr}$ and $Q_{rphi}$. (H-I) The cell elongation pattern shown in B, colored by the $Q_{rr}$ (H) or $Q_{rphi}$ (I) component.

### Fig S2, related to Fig 1: Profiles of cell elongation and dynamics around the DV boundary

#### Area and elongation in the band around the DV boundary during mid-third instar (96hr AEL)

**A. Region of interest highlighted on one frame** **B. Profile in cell area along X axis** **C. Profile in cell elongation along X axis**

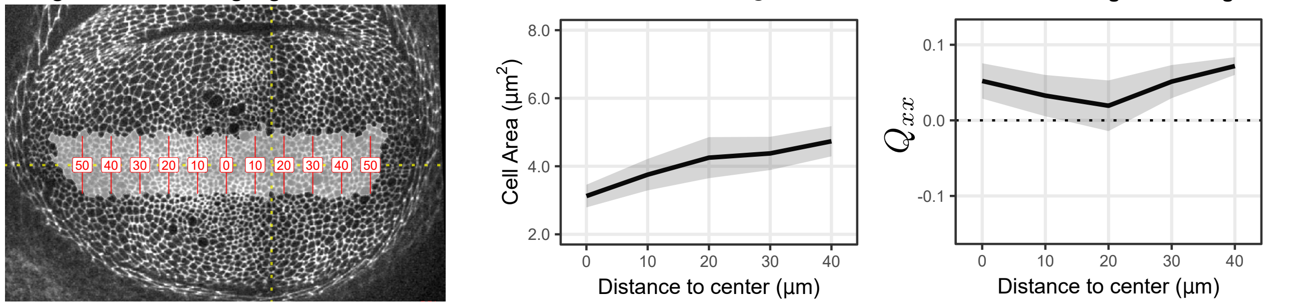


#### Gradients in growth rate in the band around the DV boundary

**D. Rate of tissue and cell area change** **E. Rate of cell divisions and extrusions**

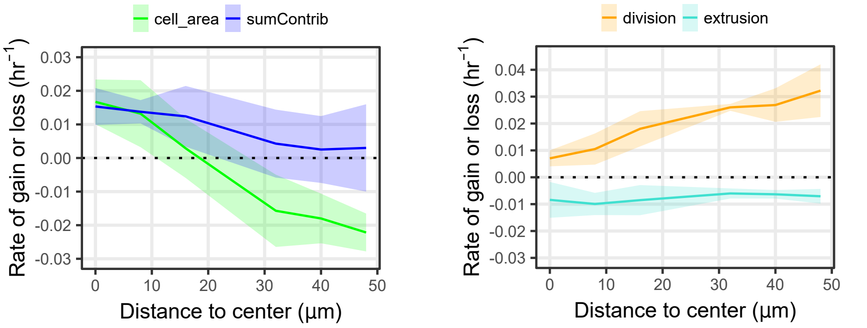


The cell area and elongation are reported for the cells surrounding the DV boundary, the region that was excluded from the analysis in Fig 1. (A) The radially elongated blade cells within the DV boundary region are shaded in grey on a single image of the timelapse of Ecadherin-GFP expressing wing discs (explanted at 96hr AEL). Red lines indicate bins along the x axis (aligned with the DV boundary) emanating from the center of elongation symmetry; the numbers indicate the distance to the center (in microns). Average cell area (B) and the XX component of cell elongation (C) were calculated by averaging over all five movies in the middle time window within the bins along the DV boundary (highlighted in A). The solid line indicates the mean, and the shaded region indicates the standard deviation. (D) The profile of tissue growth or that contributed by changes in cell area, analyzed in the bins along the DV boundary that are highlighted in A. The blue line labelled "sumContrib" is the sum of all cellular contributions, corresponding to the total tissue growth. (E) The profile along the DV boundary in the contribution of cell divisions or cell extrusions to overall tissue deformation, analyzed in the bins shown in A.

### Fig S3, related to Fig 1: Change in cell area and elongation during timelapse (96hr AEL)

#### Area and elongation in the band around the DV boundary

**A. Region of interest**

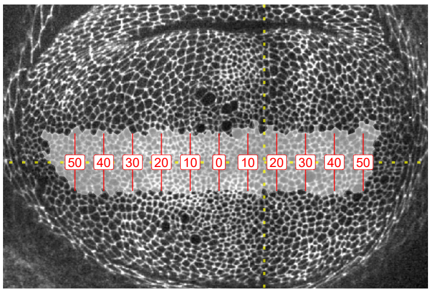


**B. Cell area and elongation profiles plotted at different intervals of the timelapse**

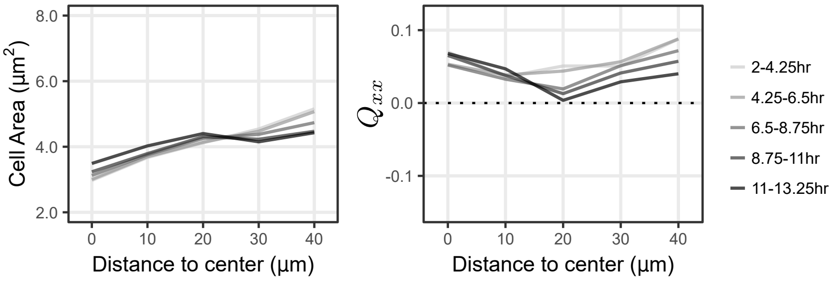


#### Area and elongation in the region outside the DV boundary band

**C. Region of interest**

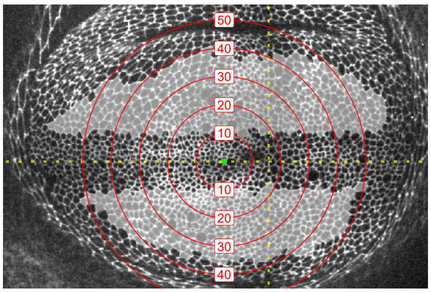


**D. Cell area and elongation profiles plotted at different intervals of the timelapse**

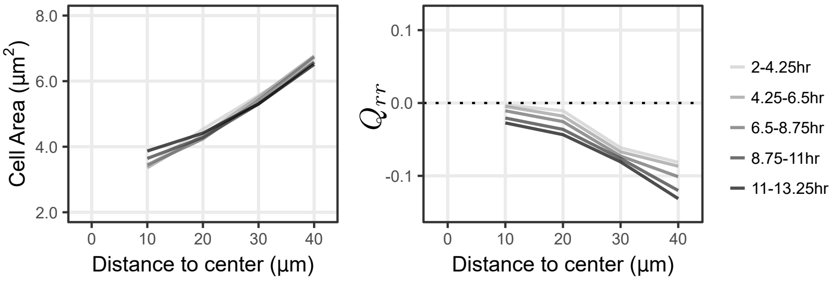


Analysis of cell area and elongation for the region of cells inside (A-B) or outside (C-D) the band around the DV boundary, analyzed over all timepoints of the movie. This analysis complements the analysis of the middle time window shown in Fig 1. (A+C) Single timepoints from the timelapse of an Ecadherin-GFP-expressing wing disc growing in *ex vivo* culture (explanted at 96hr AEL), highlighting the cells that correspond to these two regions of interest. Red lines indicate the binning from the distance to the center, either along X (A) or radially (C). Numbers indicate distance to the center (in microns). (B+D) Profiles in cell area (left) or cell elongation (right). For the band around the DV boundary (B), we present $Q_{xx}$, whereas outside this band (D), we present $Q_{rr}$. Data were averaged across 5 windows for all 5 discs, at different distances away from the center of symmetry to show how the profile changes in time during the timelapse.

### Fig S4, related to Fig 4: Analysis of circular laser ablations (96hr AEL)

#### Laser ablations in the region outside the DV boundary

A. Distribution of fit residuals | B. Anisotropic stress vs. cell elongation | C. Determining the ratio of elastic constants

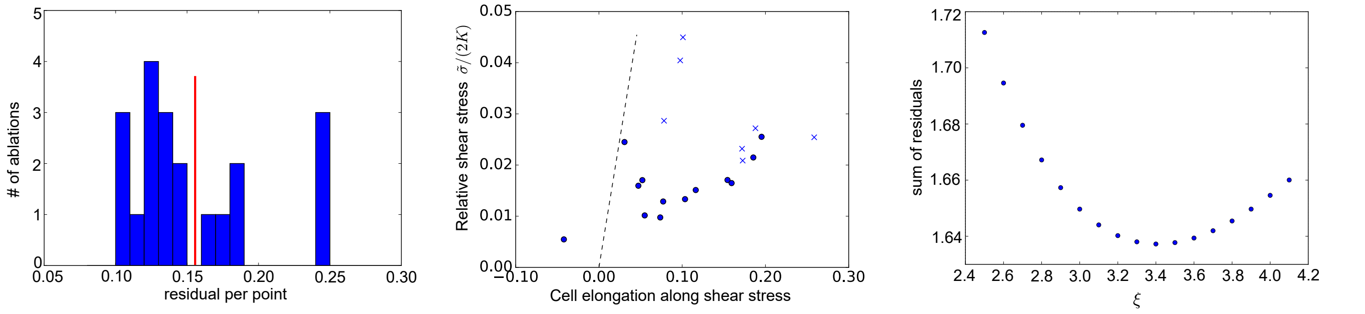


#### Laser ablations in the band around the DV boundary band

D. Destribution of fit residuals | E. Anisotropic stress vs. cell elongation | F. Determining the ratio of elastic constants

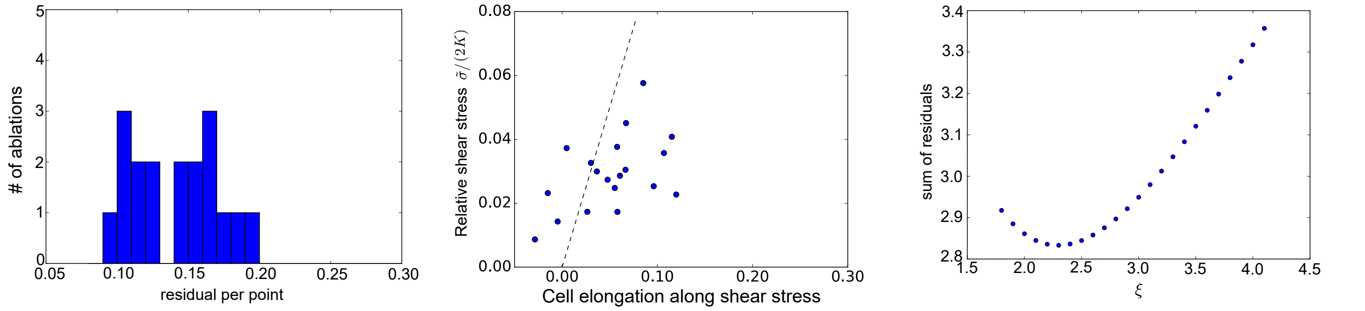


Tissue stress was related to cell elongation in mid-third instar wing discs using ESCA. (A) and (D): Histogram of fit residuals for the region outside the DV boundary (A) or within the band around the DV boundary (D). The red line in (A) denotes the cutoff we chose to discard cuts with non-elliptical outlines. (B) and (E) Plot of the anisotropic stress (relative to 2K) versus the cell elongation along the stress axis. Points marked with "x" were discarded because their fit residuals were greater than our cutoff (red line in A). Dotted line indicates a slope=1, which would correspond to a tissue without polarity-driven stress. (C) and (F) show the sum of fit residuals for a range of values for the ratio of elastic constants. The optimal value was considered to be the one that minimizes this sum of residuals.

#### Fig S5, related to Fig 7: Spatial pattern of cell elongation upon MyoVI knockdown

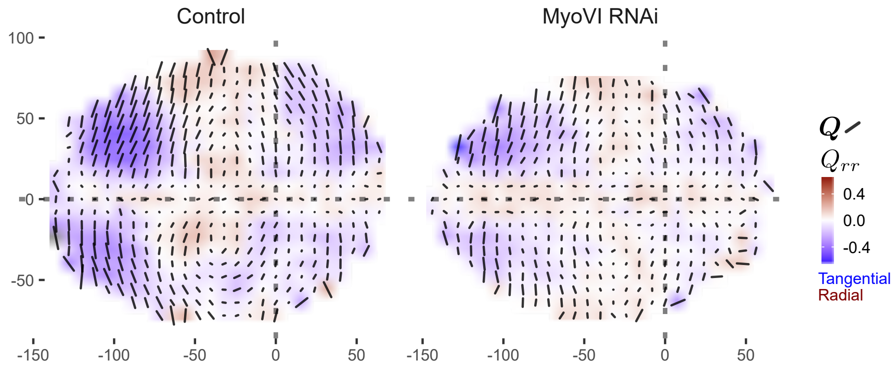


RNAi was induced against Myosin VI in the pouch (nubGAL4 > UAS-myoVIRNAi), and cell elongation was assessed at the end of larval development (119hr after egg laying), along with the corresponding control (nubGAL4/+). The cell elongation pattern was quantified in a grid centered on the AP and DV boundaries and averaged across several discs for each genotype. The bars represent the cell elongation tensor **Q**, where the length of the bar is proportional to the magnitude and the angle indicates its orientation. In addition, the radial component of cell elongation is presented as a color code, with blue indicating tangential and red indicating radial. The dotted grey lines indicate the positions of the compartment boundaries, and the axes correspond to the distance (in microns) from these boundaries.

### Supplement Theory for "Self-organized patterning of cell morphology via mechanosensitive feedback"

Natalie A. Dye,[1,2] Marko Popović,[3] K. Venkatesan Iyer,[1,2] Suzanne Eaton,[1,2] and Frank Jülicher[2,4]
[1] *Max Planck Institute for Molecular Cell Biology and Genetics, Pfotenhauerstrasse 108, 10307 Dresden, Germany*
[2] *Cluster of Excellence Physics of Life, TU Dresden, 01307 Dresden, Germany*
[3] *Institute of Physics, École Polytechnique Fédérale de Lausanne, CH-1015 Lausanne, Switzerland*
[4] *Max Planck Institute for Physics of Complex Systems, Nöthnitzer Strasse 38, 01187 Dresden, Germany*

#### CONTENTS

I. Continuum model of the wing disc epithelium 1
A. Model definition 1
B. Cell area profiles follow from cell elongation profiles 1
C. Laser ablation experiments provide an estimate of model parameters 2
Comparison of laser ablation and area profile fits 2
D. Mechanosensitivity leads to spontaneous cell polarity by self-organization 3
1. Homogeneous tissue 3
2. Polarisation of a radially symmetric tissue 3
3. Predictions for experiments 3

II. Theory of circular laser ablations 4
A. Linear elasticity in polar coordinates 4
1. Displacement gradient 4
2. Linear elastic constitutive relation 4
3. Force balance and compatibilty: Airy stress function 4
B. Circular ablation of an elastic sheet under stress 4
1. Inner piece 4
2. Outer piece 5

III. Correlation between cell elongation and cell movement contributes to the tissue radial shear flow 7

References 8

#### I. CONTINUUM MODEL OF THE WING DISC EPITHELIUM

##### A. Model definition

We use a previously developed continuum model to describe tissue mechanics [1, 2]. Tissue shear flow $\tilde{v}_{ij}$ consists of convected co-rotational change of cell elongation $Q_{ij}$ and a contribution from cell rearrangements $R_{ij}$

$$\tilde{v}_{ij} = \frac{DQ_{ij}}{Dt} + R_{ij} \quad . \tag{1}$$

Tissue shear stress consists of an elastic part and a contribution from nematic cell polarity $q_{ij}$

$$\tilde{\sigma}_{ij} = 2KQ_{ij} + \zeta q_{ij} \quad . \tag{2}$$

Shear flow due to cell rearrangements consists of term accounting for cell shape relaxation and a contribution due to nematic cell polarity $q_{ij}$

$$R_{ij} = \frac{1}{\tau}Q_{ij} + \lambda q_{ij} \quad . \tag{3}$$

Force balance couples shear stress and area pressure

$$\partial_j \tilde{\sigma}_{ij} - \partial_i P = 0 \tag{4}$$

which will be reflected in cell area. We use a constitutive relation

$$P = -\overline{K} \ln\left(\frac{a}{a_0}\right) \quad , \tag{5}$$

where $a$ represents cell area and $a_0$ corresponds to value of $a$ at $P = 0$.

We consider a simple model in which the tissue is a radially symmetric disc, see Fig. 3 C-E of the main text. We use polar coordinate system $(r, \varphi)$, where the radial force balance reads

$$\partial_r P = \partial_r \tilde{\sigma}_{rr} + \frac{2}{r}\tilde{\sigma}_{rr} \tag{6}$$

Furthermore, in the model we consider a constrained tissue with negligible radial shear flow $\tilde{v}_{rr} = 0$, consistent with experimental observations, see Fig. 2 of the main text.

##### B. Cell area profiles follow from cell elongation profiles

Empirically, the radial cell elongation profile can be described by a power law

$$Q_{rr} \simeq -\left(\frac{r}{r_Q}\right)^{\theta} \quad , \tag{7}$$

with values for $r_Q = 140 \pm 20\mu m$ and $\theta = 1.6 \pm 0.1$ obtained by fitting the data on the last $\sim 5$ hours of the

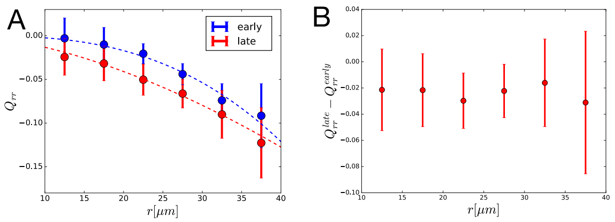


Theory Supplement Figure 1. Left: Radial cell elongation in the first half of the movies ('early') are shown in blue and in the second half of the movies ('late') are shown in red. Fits of power law are shown with dashed lines in corresponding colors, with fit parameters $r_Q = 84 \pm 3\mu m$, $\theta = 2.8 \pm 0.3$ and $r_Q = 140 \pm 20\mu m$, $\theta = 1.6 \pm 0.1$ for 'early' and 'late' profiles, respectively. Right: Radial cell elongation change suggests that there is no significant gradient of radial shear due to cell rearrangements $R_{rr}$ during the experiment.

experiments, see red line in Theory Supplement Figure 1 A. We find $r_Q = 84 \pm 3\mu m$ and $\theta = 2.8 \pm 0.3$ by fitting the data from the first $\sim 5$ hours. Data is averaged over 5 experiments and errorbars represent a standard deviation among the mean values in each experiments.

At this point it is useful to estimate the spatial profile of shear due to cell rearrangements $R_{rr}$. We calculate the difference between late and early cell elongation profiles, as shown in Theory Supplement Figure 1 B. Since $\tilde{v}_{rr} \approx 2 \cdot 10^{-3} h^{-1}$ we find $R_{rr} \approx (6 \pm 7) \cdot 10^{-3} h^{-1}$ with no significant variation in $r$.

Now, we find an expression for cell area using Eqs. 2, 3, 5, 6 and 7

$$a = a_0 e^{-\frac{c_a}{\overline{K}} + \frac{2K^*}{\overline{K}} \frac{\theta+2}{\theta} \left(\frac{r}{r_Q}\right)^{\theta} - \frac{2K}{\overline{K}} \left(1 - \frac{K^*}{K}\right) \tau R_{rr} \ln\left(\frac{r}{r_Q}\right)}, \quad (8)$$

where

$$K^* = \left(1 - \frac{\zeta}{2K\tau\lambda}\right) K \quad (9)$$

and $c_a$ is an integration constant. We fit the Eq. 8 to the experimental data as shown in Fig. 3 G of the main text, and we find parameter values

$$a_0 e^{-\frac{c_a}{\overline{K}}} = 3.3 \pm 0.3 \quad , \quad (10)$$

$$\frac{2K^*}{\overline{K}} \frac{\theta + 2}{\theta} = 5.4 \pm 0.7 \quad , \quad (11)$$

$$\frac{2K}{\overline{K}} \left(1 - \frac{K^*}{K}\right) \tau R_{rr} = 0.00 \pm 0.07 \quad , \quad (12)$$

where the reported uncertainties represent square roots of the fit covariance matrix and the parameter in Eq. 12 was constrained to be positive, motivated by the fact that we do not find $K^* > K$ (see Eq. 18 below).

#### C. Laser ablation experiments provide an estimate of model parameters

We write Eq. 2 in form

$$\frac{\tilde{\sigma}_{rr}}{2K} = \frac{K^*}{K} Q_{rr} + \left(1 - \frac{K^*}{K}\right) \tau R_{rr} \quad . \quad (13)$$

We fit this equation to the experimentally obtained shear stress as a function of cell elongation in the laser ablation experiments. We find

$$\frac{K^*}{K} = 0.05 \pm 0.02 \quad , \quad (14)$$

$$\left(1 - \frac{K^*}{K}\right) \tau R_{rr} = 0.011 \pm 0.002 \quad . \quad (15)$$

Using the estimate for $R_{rr}$ we find

$$\tau = (2 \pm 2)h \quad . \quad (16)$$

This value is consistent with the one found for the cell elongation relaxation time-scale in the pupal wing [1].

In Eq. 3 we can now show that cell elongation profile in the wing pouch reflects the profile of nematic cell polarity. Namely, $|Q_{rr}| \gg \tau |R_{rr}| \approx 0.012$ in most of the tissue, except in the vicinity of the DV boundary, and thus

$$q_{rr} \approx -\tau\lambda Q_{rr} \quad . \quad (17)$$

*Comparison of laser ablation and area profile fits*

It is interesting to notice that from Eq. **??** and the value for $2K/\overline{K} = 3.4 \pm 0.4$ obtained from the laser ablation experiments (main text and Methods), we can estimate

$$\frac{K^*}{K} = 0.6 \pm 0.1 \quad . \quad (18)$$

This value is significantly higher than the one obtained by a direct fit to the laser ablation data (Eq. 14). This apparent discrepancy stems in part from the fact that the elastic constant $\overline{K}$ in Eq. 5 is a non-linear elastic modulus. It can only be related the one used in the laser ablation method by linearizing it around a typical value of $a$ in the data. When this value is not equal to $a_0$ the linearized elastic constant will differ from the original $\overline{K}$ in Eq. 5. Furthermore, Eq. 5 does not take into account any adaptation of cell area to pressure or nematic cell polarity, which could influence the measured value $K^*/K$ when fitting the cell area profile. However interesting, these effects are beyond the scope of our current work.

#### D. Mechanosensitivity leads to spontaneous cell polarity by self-organization

##### 1. Homogeneous tissue

We propose a dynamical equation of the polarity tensor $q_{ij}$

$$\frac{Dq_{ij}}{Dt} = -\frac{1}{\tau_q} q_{ij} - \mu\tilde{\sigma}_{ij} \quad , \tag{19}$$

where the parameter $\mu$ represents the mechanosensitive response of nematic cell polarity to shear stress. We use Eqs. 1, 2, 3 and 19 to obtain a dynamic equation for shear stress

$$\begin{aligned}\frac{\partial\tilde{\sigma}_{ij}}{\partial t} &= 2K\tilde{v}_{ij} - \left(\frac{1}{\tau} + \zeta\mu\right)\tilde{\sigma}_{ij} \\ &- \left(2K\lambda + \zeta\left(\frac{1}{\tau_q} - \frac{1}{\tau}\right)\right) q_{ij} \quad .\end{aligned} \tag{20}$$

The system of linear equations 19 and 20 becomes unstable when

$$2K^*\tau_q\mu > 1 \quad . \tag{21}$$

This instability will result in a finite value of nematic cell polarity in the tissue. To account for this polarised state we have to include a stabilising higher order term in Eq. 19

$$\frac{Dq_{ij}}{Dt} = -\frac{1}{\tau_q} q_{ij} - \mu\tilde{\sigma}_{ij} - \alpha q^2 q_{ij} \quad . \tag{22}$$

In the steady state the magnitude squared of the nematic cell polarity is

$$q^2 = q_0^2 \equiv \frac{2K^*\tau_q\mu - 1}{\alpha\tau_q} \quad . \tag{23}$$

Here, we have introduced the strength of polarity $q_0$, corresponding to the magnitude of polarity in a homogeneous system. Note that the definition of $q_0^2$ allows for negative values, which would correspond to no spontaneous polarisation in the tissue.

##### 2. Polarisation of a radially symmetric tissue

To account for spatial variations of nematic cell polarity, we introduce a Laplacian term in Eq. 22

$$\frac{Dq_{ij}}{Dt} = -\frac{1}{\tau_q} q_{ij} - \mu\tilde{\sigma}_{ij} - \alpha q^2 q_{ij} + D\nabla^2 q_{ij} \tag{24}$$

To fit the radial profile of $Q_{rr}$ observed in time-lapse experiments we account for non-vanishing $R_{rr}$ as well. Consistent with the observation that during the experiments $R_{rr}$ does not vary significantly in time (Fig 2B main text) we use approximation $\tau\partial_t R_{rr} \ll R_{rr}$. Then $q_{rr}$ satisfies

$$\begin{aligned}\partial_r^2 q_{rr} + \frac{1}{r}\partial_r q_{rr} - \frac{q_{rr}}{r^2} &= \frac{\alpha}{D}\left(q_{rr}^3 - q_0^2 q_{rr}\right) \\ &+ \frac{1}{D}\left(\frac{1}{\tau\lambda} + 2K\mu\tau\right) R_{rr} \quad ,\end{aligned} \tag{25}$$

where the left hand side is the radial component of the Laplacian of $q_{ij}$ in polar coordinate system. From solution of this equation the $Q_{rr}$ profile is obtained as $Q_{rr} \approx -\tau\lambda q_{rr}$. This quantity is fitted to the experimentally measured tangential cell elongation profile by optimizing 4 tissue parameters, as well as two boundary conditions $q_{rr}(r_{in})$ and $\partial_r q_{rr}(r_{in})$ required to solve Eq. 25. Here $r_{in} = 10\mu m$ is the smallest radius at which we measure the tangential cell elongation profile in Fig. 6 C of the main text. Parameters obtained by the fit are reported in Table 1 of the main text and the fitted $Q_{rr}$ profile is shown in Fig. 6 of the main text. Parameter uncertainties were estimated as the 10 and 90 percentile intervals obtained by fitting uniformly sampled profiles of $Q_{rr}$ from the interval $Q_{rr}(r) + \Delta Q_{rr}(r)$ and $Q_{rr}(r) - \Delta Q_{rr}(r)$, see Fig 3 of the main text, with 101 realisations. Obtained uncertainty intervals are reported in the Table 1 of the main text.

##### 3. Predictions for experiments

Spontaneous cell polarity in our model depends on mechanosensitive feedback characterized by the feedback strength $\mu$. The characteristic polarity strength $q_0$ grows monotonically with $\mu$. A prediction from our model is therefore that when $\mu$ is reduced, cell elongation will be reduced because it is mediated by cell polarity. In order to test this prediction, we performed MyoVI knockdown experiments which indeed show a decrease in cell elongation.

Our qualitative prediction is rather robust and does not depend on details of the system such as the influence of compartment boundaries which disrupt the radial symmetry. However boundary conditions, which we do not know, could have a significant influence on the system. Therefore a full quantitative prediction of experiments is difficult. To show the self-consistency of our prediction, we test if the value of $q_0$ that we use to describe the data is consistent with the actual cell elongations observed. Indeed we have $q_0 \approx 0.1$, see Table 1 which predicts a typical strength of cell elongation $Q_0 = -\lambda\tau q_0 \approx -0.1$, consistent with experimentally measured values, see Fig. 6 C of the main text. This suggests that our qualitative prediction is robust given the uncertainties in the system.

#### II. THEORY OF CIRCULAR LASER ABLATIONS

We developed a method to infer the stresses in an epithelial tissue from the tissue boundary shape after a circular ablation. Here, we first briefly review linear elasticity in polar coordinates. Then, we derive equations relating shape of the inner and outer boundaries obtained by the ablation of an infinite elastic sheet in the small deformation regime. The application of the method and the obtained parameter estimates are described in Methods section.

##### A. Linear elasticity in polar coordinates

###### 1. Displacement gradient

The deformation of an elastic object is described by the displacement vector of each point on the sheet

$$u_i(x_j) = x'_i(x_j) - x_i \tag{26}$$

where $x_i$ is the intial position of a point on the elastic sheet and $x'_i$ is the position of that point after the deformation. Shape changes are described by the symmetric traceless gradient of displacement

$$u_{ij} = \frac{1}{2}\left(\partial_j u_i + \partial_i u_j\right) \quad , \tag{27}$$

which can be decomposed into isotropic part $u = u_{kk}$ and shear part $\tilde{u}_{ij} = u_{ij} - \delta_{ij}u/2$. Note that we use summation convention over repeated indices. In polar coordinates displacement gradient is given by

$$u_{rr} = \partial_r u_r \tag{28}$$

$$u_{\varphi\varphi} = \frac{1}{r}\partial_\varphi u_\varphi + \frac{1}{r}u_r \tag{29}$$

$$u_{r\varphi} = \frac{1}{2}\left(\partial_r u_\varphi - \frac{1}{r}u_\varphi + \frac{1}{r}\partial_\varphi u_r\right) \tag{30}$$

###### 2. Linear elastic constitutive relation

We are considering the linear elastic constitutive relation

$$\sigma_{ij} = 2K\tilde{u}_{ij} + \zeta q_{ij} + \overline{K}u\delta_{ij} \quad , \tag{31}$$

where $\tilde{\sigma}_{ij}$ is traceless symmetric component of shear stress, with shear elastic modulus $K$ and bulk elastic modulus $\overline{K}$. Here, we allow for the presence of active stresses due to a nematic field $q_{ij}$, corresponding to the nematic cell polarity in the wing disc. However, a constant active stress does not influence our results in the small deformation limit, since it simply redefines the reference configuration. We can therefore write this constitutive equation as

$$u_{ij} = \frac{1}{2K}\tilde{\sigma}_{ij} + \frac{1}{4\overline{K}}\sigma_{kk}\delta_{ij} \quad . \tag{32}$$

###### 3. Force balance and compatibilty: Airy stress function

Force balance imposes

$$\partial_j \sigma_{ij} = 0 \quad . \tag{33}$$

In two dimensions, force balance is satisfied by all stress fields whose components are derived from the Airy stress function $\Phi$

$$\begin{aligned} \sigma_{xx} &= \partial_y^2 \Phi \quad , \\ \sigma_{xy} &= -\partial_x \partial_y \Phi \quad , \\ \sigma_{yy} &= \partial_x^2 \Phi \quad , \end{aligned} \tag{34}$$

which can be expressed in polar coordinates

$$\sigma_{rr} = \frac{1}{r}\partial_r \Phi + \frac{1}{r^2}\partial_\varphi^2 \Phi \quad , \tag{35}$$

$$\sigma_{r\varphi} = -\partial_r\left(\frac{1}{r}\partial_\varphi \Phi\right) \quad , \tag{36}$$

$$\sigma_{\varphi\varphi} = \partial_r^2 \Phi \quad . \tag{37}$$

Now, a compatibility condition, which follows from requirement of continuity of displacement field, can be written as

$$\nabla^4 \Phi = 0 \quad . \tag{38}$$

The general solution of this equation in polar coordinates, also called Michell solution, is given by

$$\begin{aligned} \Phi = &\left[A_0 r^2 + B_0 r^2 \ln r + C_0 \ln r\right] \\ &+ \left[I_0 r^2 + I_1 r^2 \ln r + I_2 \ln r + I_3\right]\varphi \\ &+ \left[A_1 r + B_1 r^{-1} + \overline{B}_1 r\varphi + C_1 r^3 + D_1 r \ln r\right]\cos\varphi \\ &+ \left[E_1 r + F_1 r^{-1} + \overline{F}_1 r\varphi + G_1 r^3 + H_1 r \ln r\right]\sin\varphi \\ &+ \sum_{n>1}\left[A_n r^n + B_n r^{-n} + C_n r^{n+2} + D_n r^{-n+2}\right]\cos n\varphi \\ &+ \sum_{n>1}\left[E_n r^n + F_n r^{-n} + G_n r^{n+2} + H_n r^{-n+2}\right]\sin n\varphi \quad . \end{aligned} \tag{39}$$

Inferring the stresses from laser ablation experiments consists of solving two elastic problems using the Michell solution as we now show.

##### B. Circular ablation of an elastic sheet under stress

Laser ablation splits the elastic sheet into inner and outer pieces. We now show that the boundary shapes of the two pieces are ellipses by solving the corresponding elastic problems.

###### 1. Inner piece

We consider a piece of elastic sheet stretched by stress $\sigma_{xx} = \sigma^0_{xx}$, $\sigma_{yy} = \sigma^0_{yy}$, $\sigma_{xy} = 0$. After a circular laser ablation the inner piece is under no stress and to infer the original stresses in the sheet we determine the shape of the sheet boundary assuming the stresses are known. Then, given the boundary shape we will be able to infer the stresses.

Stress in a sheet that is in mechanical equilibrium, under no external body force, is uniform throughout the sheet. Therefore, using Eqs. 28 we can integrate Eq. 32 to find the components of the displacement vector $u_i$

$$u_r = \frac{r}{2K}\left[\tilde{\sigma}\cos 2\varphi + \frac{K}{2\overline{K}}\sigma\right] \tag{40}$$

$$u_\varphi = -\frac{r}{2K}\tilde{\sigma}\sin 2\varphi \quad , \tag{41}$$

where we have introduced

$$\sigma = \sigma^0_{xx} + \sigma^0_{yy} \tag{42}$$

$$\tilde{\sigma} = (\sigma^0_{xx} - \sigma^0_{yy})/2 \quad . \tag{43}$$

To relate the displacement to the shape of the inner piece boundary, we have to solve Eq. 26 for $r(\varphi)$, in the configuration illustrated in Theory Supplement Figure 2

$$r\hat{r} + u_r\hat{r} + u_\varphi\hat{\varphi} = R\hat{r}' \quad . \tag{44}$$

Using the fact that

$$\hat{r}'\cdot\hat{r} = \cos(\varphi' - \varphi) \tag{45}$$

$$\hat{r}'\cdot\hat{\varphi} = \sin(\varphi' - \varphi) \tag{46}$$

we obtain

$$r\left(\varphi\middle|\frac{\sigma}{\overline{K}}, \frac{\tilde{\sigma}}{K}\right) = \frac{R}{\sqrt{\left[1 + \frac{\sigma}{4\overline{K}} + \frac{\tilde{\sigma}}{2K}\cos(2\varphi)\right]^2 + \left[\frac{\tilde{\sigma}}{2K}\sin(2\varphi)\right]^2}} \tag{47}$$

To simplify the notation we introduce

$$\tilde{s} = \frac{\tilde{\sigma}}{2K} \tag{48}$$

$$s = \frac{\sigma}{\overline{K}} \quad , \tag{49}$$

and we write the solution in the form

$$r(\varphi|s,\tilde{s}) = \frac{R}{\sqrt{\left[1 + \frac{1}{4}s + \tilde{s}\cos(2\varphi)\right]^2 + \left[\tilde{s}\sin(2\varphi)\right]^2}} \quad . \tag{50}$$

We denote semiaxes of this ellipse along the $x$ and $y$ axes, as defined in Theory Supplement Figure 2, by $a_{\text{in}}$ and $b_{\text{in}}$, respectively. We find that

$$a_{\text{in}} = \frac{R}{|1 + \frac{1}{4}s + \tilde{s}|} \tag{51}$$

$$b_{\text{in}} = \frac{R}{|1 + \frac{1}{4}s - \tilde{s}|} \tag{52}$$

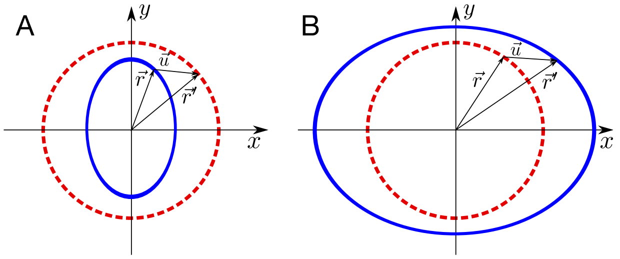


Theory Supplement Figure 2. A. Inner piece problem: Blue ellipse shape of the boundary after the ablation is the reference shape deformed by uniform stress $\sigma_{xx} = \sigma^0_{xx}$, $\sigma_{yy} = \sigma^0_{yy}$ to the red circle boundary shape before the ablation. B) Outer piece problem: The red circle boundary shape before the ablation is the reference in this case. The final boundary shape after the ablation is blue ellipse.

##### 2. *Outer piece*

As in the case of the inner piece we need to find the boundary shape as a function of the original stresses in the sheet. In this case, the normal stress at the ablation boundary is zero, but the sheet is not stress-free. To simplify the problem, we consider initially stress-free sheet with a circular hole to which boundary stresses $-\sigma^0_{ij}$ are applied. For small deformations, where elastic problem is linear and solutions of the problems commute, the boundary shape is equivalent to the one created by the laser ablation of a sheet under stress $\sigma^0_{ij}$.

Now we have solve the full elastic problem since stresses are not uniform. We keep terms of the Airy stress function $\Phi$, which obey nematic symmetry of the problem and relax sufficiently fast at infinity. We also assume that the polar axis is oriented along one of the principal axes of stress tensor such that the off-diagonal component of stress vanishes. We obtain

$$\Phi = C_0\ln r + \left(D_2 + \frac{B_2}{r^2}\right)\cos(2\varphi) \tag{53}$$

and for stress components, using Eq. 35

$$\sigma_{rr} = \frac{C_0}{r^2} - 2\left(2\frac{D_2}{r^2} + 3\frac{B_2}{r^4}\right)\cos(2\varphi) \tag{54}$$

$$\sigma_{r\varphi} = -2\left(\frac{D_2}{r^2} + 3\frac{B_2}{r^4}\right)\sin(2\varphi) \tag{55}$$

$$\sigma_{\varphi\varphi} = -\frac{C_0}{r^2} + 6\frac{B_2}{r^4}\cos(2\varphi) \tag{56}$$

The stress boundary condition reads

$$\sigma_{rr}(r=R) = -\frac{\sigma}{2} - \tilde{\sigma}\cos(2\varphi) \tag{57}$$

$$\sigma_{r\varphi}(r=R) = \tilde{\sigma}\sin(2\varphi) \tag{58}$$

allows us to determine coefficients in Eq. 54

$$C_0 = -\frac{1}{2}\sigma R^2 \tag{59}$$

$$B_2 = -\frac{1}{2}\tilde{\sigma} R^4 \tag{60}$$

$$D_2 = \tilde{\sigma} R^2 \quad . \tag{61}$$

Now we can calculate and then integrate the deformation gradient components to obtain

$$u_r = R\left[\frac{\overline{K}}{4K}\frac{\sigma}{\overline{\overline{K}}}\frac{R}{r} + \left[2\left(1+\frac{K}{\overline{\overline{K}}}\right)\frac{R}{r} - \frac{R^3}{r^3}\right]\frac{\tilde{\sigma}}{2K}\cos(2\varphi)\right] \tag{62}$$

$$u_\varphi = -R\left[2\frac{K}{\overline{\overline{K}}}\frac{R}{r} + \frac{R^3}{r^3}\right]\frac{\tilde{\sigma}}{2K}\sin(2\varphi) \quad . \tag{63}$$

Finally, as for the inner cut, we have to solve

$$R\hat{r} + u_r\hat{r} + u_\varphi\hat{\varphi} = r'\hat{r}' \tag{64}$$

for $r'(\varphi')$. We express the solution using the angle $\varphi$ as a parameter

$$\varphi'\left(\varphi\middle|\frac{\sigma}{\overline{\overline{K}}}, \frac{\tilde{\sigma}}{2K}, \frac{K}{\overline{\overline{K}}}\right) = \varphi - \arctan\left(\frac{\left(1+2\frac{K}{\overline{\overline{K}}}\right)\frac{\tilde{\sigma}}{2K}\sin(2\varphi)}{1+\frac{\overline{K}}{4K}\frac{\sigma}{\overline{\overline{K}}} + \left(1+2\frac{K}{\overline{\overline{K}}}\right)\frac{\tilde{\sigma}}{2K}\cos(2\varphi)}\right) \tag{65}$$

$$r'\left(\varphi\middle|\frac{\sigma}{\overline{\overline{K}}}, \frac{\tilde{\sigma}}{2K}, \frac{K}{\overline{\overline{K}}}\right) = R\sqrt{\left[1+\frac{\overline{K}}{4K}\frac{\sigma}{\overline{\overline{K}}} + \left(1+2\frac{K}{\overline{\overline{K}}}\right)\frac{\tilde{\sigma}}{2K}\cos(2\varphi)\right]^2 + \left[\left(1+2\frac{K}{\overline{\overline{K}}}\right)\frac{\tilde{\sigma}}{2K}\sin(2\varphi)\right]^2} \quad . \tag{66}$$

Note ratio of elastic constants appears as a parameter of shape.

Now we show that the shape defined by Eqs. 65 and 66 is an ellipse. To this end, we first define

$$A = \left(1+\frac{2K}{\overline{\overline{K}}}\right)\frac{\tilde{\sigma}}{2K} \tag{67}$$

$$B = 1+\frac{\overline{K}}{4K}\frac{\sigma}{\overline{\overline{K}}} \quad . \tag{68}$$

This allows us to write Eqs. 65 and 66 as

$$\varphi'(\varphi) = \varphi - \arctan\left(\frac{A\sin(2\varphi)}{B+A\cos(2\varphi)}\right) \tag{69}$$

$$r'(\varphi) = R\sqrt{A^2+B^2+2AB\cos 2\varphi} \quad . \tag{70}$$

Now we show that $r'(\varphi')$ is an ellpise. An ellipse equation can be written as

$$r'(\varphi') = \frac{a_{\rm out}b_{\rm out}}{\sqrt{b_{\rm out}^2\cos^2\varphi' + a_{\rm out}^2\sin^2\varphi'}} \quad , \tag{71}$$

where $a_{\rm out}$ and $b_{\rm out}$ are semiaxes of the ellipse oriented along $x$ and $y$ axes, respectively. Starting from Eq. 71 and expressing $\varphi'$ from Eq. 69 we will show that we can recover Eq. 70 for a particular choice of parameters $a$ and $b$.

We first calculate

$$\cos\varphi' = \frac{(A+B)\cos\varphi}{\sqrt{A^2+B^2+2AB\cos 2\varphi}} \tag{72}$$

$$\sin\varphi' = \frac{(B-A)\sin\varphi}{\sqrt{A^2+B^2+2AB\cos 2\varphi}} \quad . \tag{73}$$

Inserting these identities into Eq. 71 we find

$$r'(\varphi') = \frac{a_{\rm out}b_{\rm out}\sqrt{A^2+B^2+2AB\cos 2\varphi}}{\sqrt{b_{\rm out}^2(A+B)^2\cos^2\varphi + a_{\rm out}^2(B-A)^2\sin^2\varphi}} \quad . \tag{74}$$

Now, we only need to find $a_{\rm out}$ and $b_{\rm out}$ for which

$$\begin{aligned}\frac{a_{\rm out}^2 b_{\rm out}^2}{R^2} &= b_{\rm out}^2(A+B)^2\cos^2\varphi \\ &\quad + a_{\rm out}^2(B-A)^2\left(1-\cos^2\varphi\right) \quad ,\end{aligned} \tag{75}$$

for all $\varphi$. The solutions are

$$a_{\rm out} = R\,|A+B| \tag{76}$$

$$b_{\rm out} = R\,|B-A| \quad , \tag{77}$$

or in terms of the original parameters

$$a_{\rm out} = R\left|1+\frac{\overline{K}}{4K}\frac{\sigma}{\overline{\overline{K}}} + \left(1+\frac{2K}{\overline{\overline{K}}}\right)\frac{\tilde{\sigma}}{2K}\right| \tag{78}$$

$$b_{\rm out} = R\left|1+\frac{\overline{K}}{4K}\frac{\sigma}{\overline{\overline{K}}} - \left(1+\frac{2K}{\overline{\overline{K}}}\right)\frac{\tilde{\sigma}}{2K}\right| \quad . \tag{79}$$

If, in addition to Eqs. 48 and 49 we define

$$\xi = \frac{2K}{\overline{\overline{K}}} \quad , \tag{80}$$

we can write

$$a_{\rm out} = R\left|1+\frac{1}{2\xi}s + (1+\xi)\,\tilde{s}\right| \tag{81}$$

$$b_{\rm out} = R\left|1+\frac{1}{2\xi}s - (1+\xi)\,\tilde{s}\right| \tag{82}$$

Equations 51, 52, 81 and 82 impose 4 constraints on 3 parameters: $s$, $\tilde{s}$ and $\xi$. When determining these parameters from experimental data we are in general not be able to satisfy all 4 equations simultaneously. Instead, we find the 3 parameters that best fit the segmented outlines of tissue boundaries, see Methods.

#### III. CORRELATION BETWEEN CELL ELONGATION AND CELL MOVEMENT CONTRIBUTES TO THE TISSUE RADIAL SHEAR FLOW

In this work we use the previously developed method to define and calculate cell elongation tensor and tissue shear due to T1 transitions, cell divisions and extrusions [1, 3]. However, we apply the method in the polar coordinate system and to make it fully consistent we have to introduce a new correlation term $S_{ij}$, which arises in the polar coordinate system when translation and elongation are correlated. To demonstrate the origin and derive an expression for $S_{ij}$ we consider a tissue that translates over the time interval $\Delta t$ but does not change otherwise. The tissue is triangulated by connecting the neighboring cell centers (see [3]). Initially, the angle of a triangle $k$ with with respect to a center point is $\varphi$, blue triangle in Fig. Theory Supplement Figure 3. During the translation by $\Delta\vec{r} = \vec{r}(t+\Delta t) - \vec{r}(t)$ the radial angle of the triangles changes by a small angle $\Delta\varphi^k$. Therefore, the components of triangle elongation tensor $\tilde{q}^t_{ij}$ in polar coordinate system change by

$$\Delta\tilde{q}^t_{ij} = -2\epsilon_{ik}\tilde{q}^t_{kj}\Delta\varphi \quad . \tag{83}$$

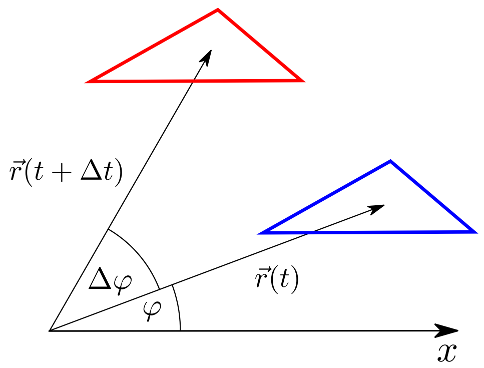


Theory Supplement Figure 3. Triangle is translated by $\Delta\vec{r} = \vec{r}(t+\Delta t) - \vec{r}(t)$. Although the triangle elongation did not change during the translation, the radial component of the triangle elongation changes due to the change of radial direction by $\Delta\varphi$.

However, since the triangles are only translating, there is no shear flow. Therefore, the average radial shear flow also vanishes and an additional term is necessary to account for the change in the average radial cell elongation in a translating tissue

$$\tilde{v}_{ij} = 0 = \frac{\Delta Q_{ij}}{\Delta t} + S_{ij} \quad . \tag{84}$$

Therefore, the correlation term is given by

$$S_{ij} = \frac{1}{\Delta t}\left\langle 2\epsilon_{ik}\tilde{q}^t_{kj}\Delta\varphi\right\rangle \quad . \tag{85}$$

In Theory Supplement Figure 4 we show the new correlation term in black measured in the pouch of the wing disc, averaged over 5 time-lapse experiments. For comparison we show the correlation term stemming from correlated fluctuations of triangle rotation and elongation in blue, and correlation term stemming from correlated fluctuations of triangle area and elongation in green. For definitions and discussion of the other two correlation terms see [1, 3]. We find that the contribution of the new correlation term to the radial shear flow is much smaller that the other two correlation terms. Therefore, contribution to the radial tissue shear flow of correlation between translation and triangle elongation is a negligible in the wing disc.

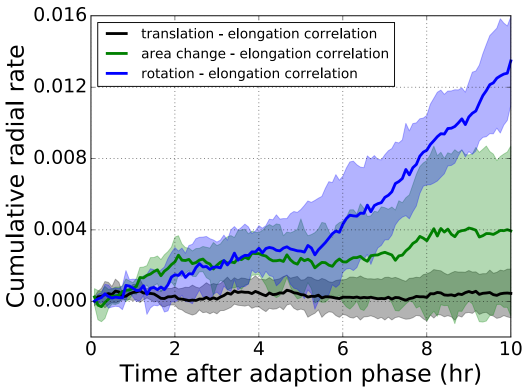


Theory Supplement Figure 4. The three correlation terms contributing to the radial tissue shear flow measured in wing disc pouch, averaged over 5 time-lapses. Shaded regions correspond to one standard deviation of that sample. Blue: Correlation term stemming from correlated fluctuations of rotation and elongation of triangles. Green: Correlation term stemming from correlated fluctuations of area and elongation of triangles. Black: The new correlation term stemming from translation and elongation of triangles. The new correlation term (black) is very small in comparison to the other two.

[1] R. Etournay, M. Popović, M. Merkel, A. Nandi, C. Blasse, B. Aigouy, H. Brandl, G. Myers, G. Salbreux, F. Jülicher, and et al., eLife **4** (2015), 10.7554/eLife.07090.
[2] M. Popović, A. Nandi, M. Merkel, R. Etournay, S. Eaton, F. Jülicher, and G. Salbreux, New Journal of Physics **19**, 033006 (2017).
[3] M. Merkel, R. Etournay, M. Popović, G. Salbreux, S. Eaton, and F. Jülicher, Physical Review E **95** (2017), 10.1103/PhysRevE.95.032401.
